# Integrative Assessment of Pathogenic Bacterial Genomes: Insights from Quality Metrics

**DOI:** 10.1101/2025.03.03.641223

**Authors:** Adeel Farooq, Asma Rafique

**Affiliations:** Research Institute for Basic Sciences (RIBS), Jeju National University, 102 Jejudaehak-ro, Jeju, 63243, Republic of Korea; Department of Microbiology and Immunology, College of Medicine, and Jeju Research Center for Natural Medicine, Jeju National University, Jeju 63243, Republic of Korea

**Keywords:** Completeness, Contiguity, Accuracy, QUAST, BUSCO, ALE

## Abstract

Genomic resources are increasingly being shared in public databases for reusability. Ensuring the quality of these resources is crucial, as the presence of low-quality genomes can compromise downstream analyses and propagate across databases. However, there is a lack of consistent and comprehensive assessments for such genomes in public genomic databases. This study focused on the quality assessment of 474 pathogenic bacterial genomes submitted from South Korea, with a particular emphasis on the integrative application of completeness, contiguity, and accuracy metrics. The genomes were ranked based on cumulative scores derived from individual metrics. High-ranked genomes consistently exceeded defined thresholds, including 98% completeness for complete single-copy BUSCO, less than 200 contigs, N50 values greater than 65 kb, and unmapped reads below 2%. Notably, positive and negative linear associations were observed among these quality metrics, emphasizing their integrative utilization, although the consistency across all metrics varied. The findings emphasize the importance of conducting comprehensive genome assessments and implementing stringent quality control measures in public databases to enhance the reliability and usefulness of pathogenic bacterial genomic data. By addressing these issues, we can improve the overall quality of public genomic resources and ensure their suitability for diverse downstream applications.

## Introduction

Scientists heavily rely on the reliability of genomic resources as they often reuse sequence data generated by others. However, if the quality of the available data is low, it not only affects downstream analysis but also propagates further into the public resources. Consequently, ensuring the quality of public data in molecular biology becomes crucial and presents a challenge for developing automated error detection and processing approaches [1].

Whole genome sequencing (WGS) has revolutionized the surveillance of human pathogenic bacteria, and numerous databases host WGS data. As of March 2021, the genomes of thirty-two significant pathogens are stored in the National Center for Biotechnology Information Pathogen Detection Database (NCBI-PD), ensuring convenient accessibility to these genomes [2]. However, strict scrutiny procedures to identify and filter out low-quality or contaminated raw sequences, misassembled genomes, and incorrectly assigned taxonomies are lacking during data submission. This poses challenges in correctly assembling raw reads, assigning accurate species taxonomies, and characterizing genes. Particularly, raw data submitted to the Sequence Read Archive (SRA) undergoes limited scrutiny at present [3].

Genome sequences often encounter quality issues such as sequencing errors, misassembly, and contamination. Contamination-related errors are particularly concerning as they can lead to misinterpretation of data, mischaracterization of gene content, inaccurate species assignment, and biases in genomic analyses. Contamination is suspected to be widespread, originating from foreign DNA present in raw biological materials or introduced during library preparation. Sequencing errors and misassembly can result in fragmented genomes, which hinder downstream analysis, including the detection of antibiotic resistance genes and mobile genetic elements such as plasmids and phages [4].

Three widely used quality control (QC) parameters for genome assessment are completeness, contiguity, and accuracy. Completeness is determined by the gene content of contigs. As contiguous genome that contains few gene contents is not useful for downstream analysis [5]. High quality genomes exhibit completeness above 90% [6]. Contiguity relates to the size and number of contigs, with high-quality genomes characterized by fewer contigs of larger size [7]. Accuracy is typically evaluated by comparing the assembled genome with a reference or by mapping reads to the assembled genome to assess coverage. Accurate genomes show consistent contig order (for reference-based assembly) or a lower percentage of unmapped reads (for de novo assembly [8].

Individual measures of contiguity, completeness, or accuracy alone can be misleading when judging genome quality [7]. Although numerous quality control measurements are available, there is currently no consistent framework for integrating these metrics to accurately report the quality of a whole genome [9]. Hence, an integration of quality control parameters is necessary for precise genome quality assessment.

Given the importance of reliable genomic resources and the potential drawbacks of low-quality genomes in public databases, it is crucial to address the gaps in quality assessment and control. Present study aimed at assessing the quality of 474 pathogenic bacterial genomes submitted from South Korea, by evaluating completeness, contiguity, and accuracy metrics. We hypothesized that integrating multiple quality metrics would provide a more comprehensive evaluation of genome quality. Our findings will contribute to improving the overall quality and utility of genomic resources in the field of molecular biology.

## Material and Methods

### Data Retrieval

We selected 474 pathogenic bacterial genomes originating from South Korea available in the NCBI-PD database. The selection criteria included the availability of SRA accession number, platform information, and raw reads. These genomes represented 13 different species (**Supplementary file 1**). Raw reads generated from Illumina-based platforms were retrieved from SRA using the SRA Toolkit (http://ncbi.github.io/sra-tools/). Briefly, prefetch was used to fetch the SRA accession numbers, and the validity of the raw sequencing data was determined using vdb-validate. Forward and reverse reads were retrieved using fastq-dump with the --split-file parameter.

### Quality Assessment of Whole Genomes

Quality trimming of the raw reads was performed using fastp [10]. Genomes were assembled using SPAdes assembler [11] with default settings, and subsequent polishing of the assembled genomes was carried out using Pilon [12]. The taxonomy of the genomes was determined through 16S RDP classifier [13]. To assess genome quality, we employed the benchmarking universal single copy orthologs (BUSCO) tool [14] with the --auto-lineage-prok detection parameter. BUSCO measures several metrics, including complete single copy, duplicated, fragmented, and missing orthologs in the genomes. The contiguity of the assembled genomes was evaluated using QUAST, a quality assessment tool [15], with default parameters. Contiguity features including number of contigs, and N50 values (defined as the length of the contig at which half of the genome is represented by contigs of that size or larger [7]), were determined. Accuracy which examines the positions of sequence read pairs within an assembly to identify anomalies [8], was evaluated using the Assembly Likelihood Evaluation (ALE) framework [16]. ALE measures the ratio of mapped and unmapped reads to the assembled genome. The complete workflow for genome quality assessment is illustrated in **Figure S1**.

### Statistical Analysis

Correlations between parameters were analyzed using Spearman’s rank order test, with a significance level set at p < 0.05. Z-scores were calculated for each parameter individually, and then the individual scores were summed. This approach allowed us to reward or penalize genomes based on their performance across multiple metrics. The quality levels, high and low, were assigned to each genome based on the standard deviation from the median value. To ensure consistency in the quality assessment, z-scores for the number of contigs and percentage of unmapped reads were multiplied by -1 before summing them up, as their lower values indicated better quality genomes [17]. Additionally, the associations among the quality metrics of high-ranked genomes were evaluated by linear regression analysis using ggplot2 package in R software.

## Results and Discussion

### Taxonomy Assignment

We used the 16S RDP classifier to assign taxonomy to 474 whole genomes. Our analysis revealed that 14 of these genomes showed 16S rRNA gene contamination of other bacteria, primarily from *Bradyrhizobium* sp. (**Table S1**). Such contamination can result in incorrect taxonomy assignment and strain identification. The existence of contaminated genomes within public resources undermines the reliability of downstream analysis [18]. Notably, all of the contaminated genomes were found to be of low quality in terms of completeness, contiguity, and accuracy. This observation aligns with a previous study that reported similar contamination issues in genomes sequenced using Illumina-based short read sequencers and among publicly available genomes in NCBI [19, 20]. Therefore, implementing proper screening procedures during genome submission to public databases is crucial to improve the overall quality of these databases.

### Evaluating Completeness, Contiguity, and Accuracy

We assessed the quality of the NCBI-PD genomes based on completeness, contiguity, and accuracy parameters. The median value of complete single-copy BUSCO was 99.10% with an interquartile range (IQR) of 4.5%. The median values for duplicated, fragmented, and missing BUSCO were 0.3%, 0.10%, and 0.40%, respectively, with respective IQRs of 0.2%, 0.8%, and 0.9%. Similarly, the median values for the number of contigs, N50, and unmapped reads were 113, 112932 bp, and 0.65%, respectively, with IQRs of 209, 156297 bp, and 1.88% (**Figure 1A**). The higher IQRs of these parameters can be attributed to the presence of contaminated and low quality genomes in the dataset.

**Figure 1.**
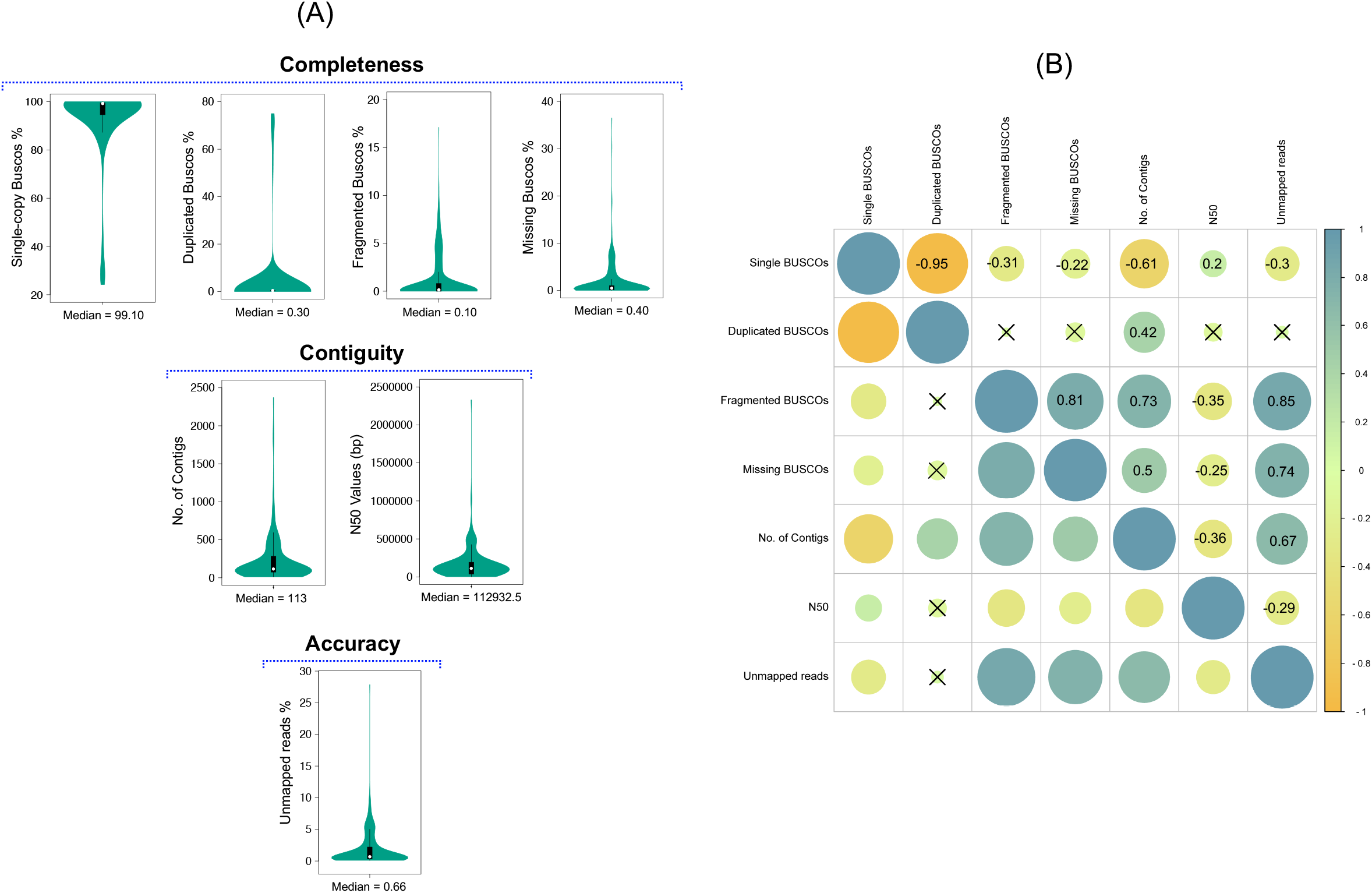
Overview of quality metrics and associations in 474 pathogenic bacterial genomes. **(A)** Violin plots illustrating the median values and interquartile ranges for seven quality parameters: single copy BUSCO, duplicated BUSCO, fragmented BUSCO, missing BUSCO, Number of contigs, N50 values, and unmapped reads. **(B)** Spearman rank order correlation analysis of genome completeness, contiguity, and accuracy metrics (n=7) using the NCBI-PD whole genomes dataset. The size and color of the circles represent the strength of the correlation. Dark blue indicates highly positive significant correlations, while dark orange indicates highly negative significant correlations. A p value < 0.05 was considered statistically significant.

Previous studies have indicated that high quality genomes typically exhibit complete single-copy BUSCO values above 90% and lower percentages of missing, duplicated, and fragmented BUSCO [21]. Consistent with this, our analysis revealed that 392 (83%) genomes had complete single-copy BUSCO values above 90%, while 444 (94%), 370 (78%), and 372 (78%) genomes had less than 2% levels of duplicated, fragmented, and missing BUSCO, respectively (**Supplementary file 2**). Well-assembled genomes from Illumina-based short-read sequencing data often exhibit a lower number of contigs and higher N50 values, which approximate the total genome size. In our study, 354 (75%) genomes had N50 values greater than 50 kb, and 346 (73%) genomes exhibited a relatively low number of contigs, with less than 200 contigs per genome. To evaluate accuracy, we measured the percentage of unmapped reads in the assembled genomes, and 348 (73%) genomes had unmapped reads below 2% (Supplementary file 2). Since more than 70% of the genomes demonstrated favorable assessment features, although not consistently across all quality metrics and with variations observed among the genomes, we proceeded to analyze the correlation among these parameters.

### Correlation Among the Assessment Parameters

The correlation among the completeness, contiguity, and accuracy parameters was assessed using Spearman’s rank order correlation coefficient. Significantly very strong, strong, moderate, and less strong correlations between the features are shown in **Figure 1B** (P value < 0.05). The percentage of complete single-copy BUSCO manifested a strong negative correlation (ρ = -0.61) with the number of contigs. Additionally, it exhibited a less strong negative correlation with unmapped reads (ρ = -0.3), and a less strong positive correlation with N50 values (ρ = 0.2). However, it is challenging to establish a consistent relationship between complete single-copy BUSCO, N50 values, and the number of contigs due to the presence of genomes with diverse quality metrics.

Some genomes with low N50 values and a higher number of contigs still had a wide range of complete single-copy BUSCO percentages, and vice versa.

We observed strong positive associations (0.73) between fragmented BUSCO and the number of contigs. Additionally, fragmented BUSCO showed very strong associations (0.8) with N50 values and unmapped reads. The percentage of missing BUSCO had a less strong negative correlation (−0.25) with N50 values, a moderate positive correlation (0.5) with the number of contigs, and a strong positive correlation (0.74) with unmapped reads. On the other hand, duplicated BUSCO values did not show a significant correlation with N50 values or unmapped reads, however they exhibited a moderate positive correlation (0.42) with the number of contigs. Although these trends were not consistent across all genomes, they provide insights into potential associations among the parameters. The negative relationship between single copy BUSCO and the number of contigs, along with the positive relationships of the other three completeness parameters, suggests that genomes with lower numbers of contigs are better assembled and exhibit higher completeness. As a result, N50 values tend to increase, and the percentage of unmapped reads decreases, as indicated by their associations. Higher percentages of fragmented and duplicated BUSCO indicate contamination in the genome sequences, often associated with an elevated number of contigs and lower N50 values. Conversely, lower percentages of these completeness parameters suggest better contiguity and accuracy of the genomes [22].

In addition, we observed a relatively less strong negative correlation (−0.36) among the contiguity features, indicating that a smaller number of contigs corresponds to higher N50 values. This relationship is logical since a smaller number of contigs represents longer contig lengths, leading to higher N50 values. Genomes with a larger number of contigs can pose challenges in downstream analysis, especially in comparative genomics tasks like gene characterization, and plasmid contig identification [23]. Better quality genomes are typically characterized by a smaller number of contigs [24], but genomes assembled from short-read sequencing methods often consist of a higher number of short contigs [25]. Therefore, in this context, genomes with a relatively smaller number of contigs tend to exhibit better N50 values compared to those with a large number of contigs [26]. Furthermore, the number of contigs showed a strong positive association (0.67) with unmapped reads, while N50 values displayed a relatively less strong negative association (− 0.29) with unmapped reads. This demonstrates the interdependence of contiguity and accuracy parameters in generating more effectively assembled genomes.

Among the BUSCO features, a strong negative association was observed between single-copy BUSCO and duplicated BUSCO, while the association of single-copy BUSCO with fragmented BUSCO and missing BUSCO was relatively less strong. Additionally, fragmented, and missing BUSCO showed a strong positive association, while duplicated BUSCO had no significant association with either of them. Since the percentage of complete single-copy BUSCO is strongly or weakly associated with the percentage of fragmented BUSCO, duplicated BUSCO, and missing BUSCO, it was chosen to represent all three features for completeness.

Although correlations were observed among the assessment metrics, we sought to derive a consistent pattern of relationships between these metrics by considering their integrative usage.

### Integrative Quality Metrics for Genomes

Correlational analysis revealed that relying on a single evaluation metric is insufficient for ranking genome quality. Thus, to effectively summarize and compare genomes, we calculated z-scores for each of the assessment feature [17]. Furthermore, we determined the rank of each genome by summing the z-scores of all metrics (Supplementary file 2). Based on the summed z-scores, we found that 237 (50%) genomes were of high rank, indicating favorable completeness, contiguity, and accuracy metrics. This finding aligns with a previous study emphasizing the use of integrative quality measures to accurately assess genome quality, as individual metrics may lead to misleading conclusions [27].

The high-ranked genomes met the established standards for quality assessment, with all of them displaying complete single-copy BUSCO values above 98%, fewer than 200 contigs, N50 values exceeding 65 kb, and unmapped reads below 2% [6, 7, 8]. These values align with recognized benchmarks for each quality metric. Additionally, an analysis of the high-ranked genomes revealed both positive and negative linear associations among the four quality metrics, highlighting their integrative usage (**Figure 2)**. Specifically, we identified a negative linear association between the percentage of single-copy BUSCO and both the number of contigs (r = -0.6), and the percentage of unmapped reads (r = -0.3). This suggests that as the percentage of complete single-copy BUSCO increases, the number of contigs and the percentage of unmapped reads tend to decrease. Conversely, we found a positive linear association between the percentage of complete single-copy BUSCO and N50 values (r = +0.2), indicating that as the percentage of complete single-copy BUSCO increases, the N50 values tend to increase. This implies that the completeness of the whole genomes, as assessed by the percentage of complete single-copy BUSCO, reflects the overall assembly quality of the genomes, although the associations may not hold consistently across all genomes [28].

**Figure 2.**
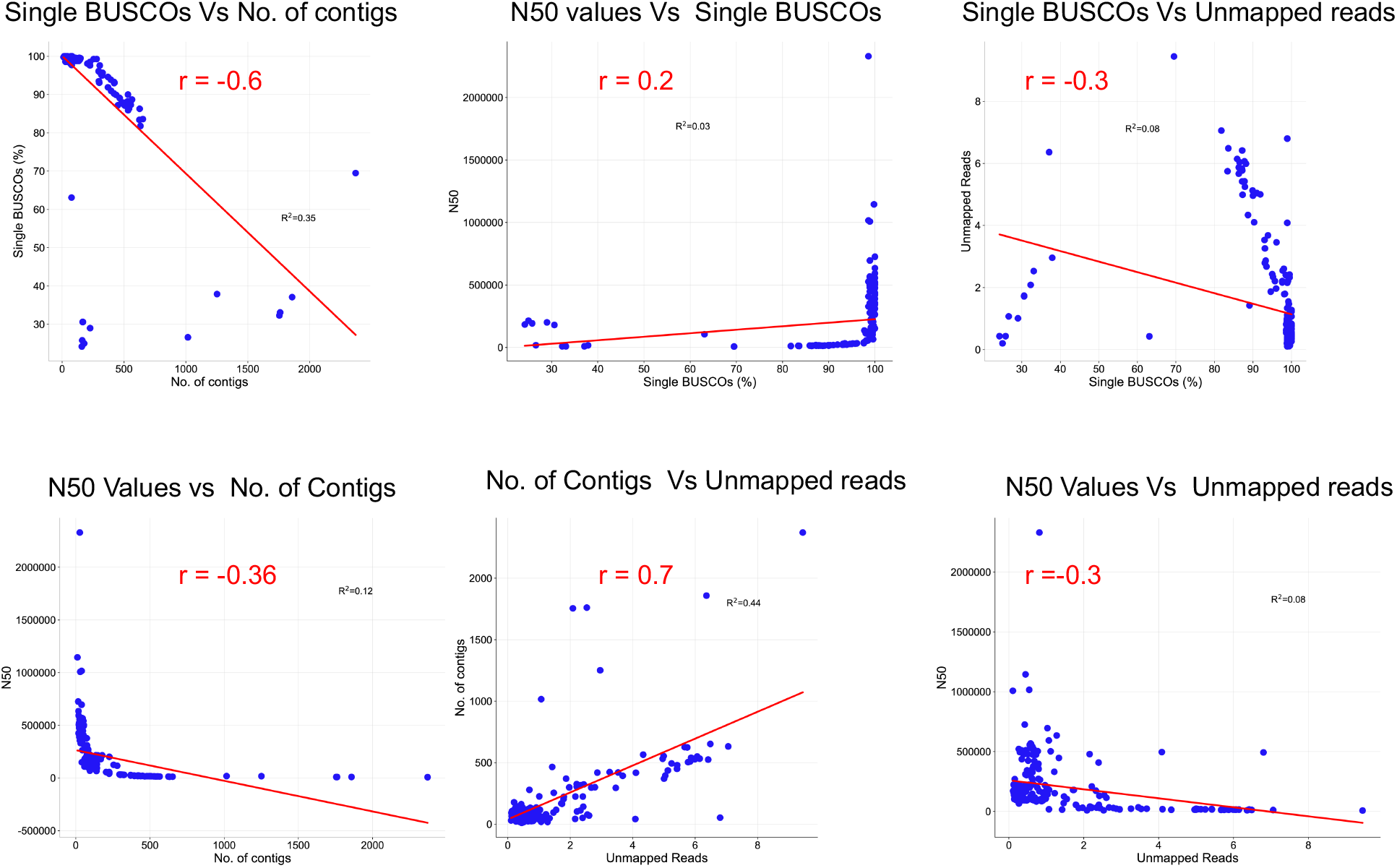
Associations between four quality metrics of completeness, contiguity, and accuracy in 237 high-ranked whole genomes. The solid line represents the linear prediction. Six distinctive graphs representing linear regression among complete single-copy BUSCO, number of contigs, N50 values, and unmapped reads.

Moreover, we found that the number of contigs exhibited a negative linear association with N50 (r = -0.4) and a positive association with unmapped reads (r = +0.7), while a negative linear relationship was also observed between N50 values and unmapped reads (r = -0.3). One of the features considered indicative of genomes quality is the maximum N50 value, which corresponds to a lower number of contigs [23]. Hence, these associations observed in the high-ranked genomes suggest the integrative usage of these metrics for precise quality assessment of pathogenic bacterial whole genomes.

It is recommended that genomic data should be validated for quality before uploading to the public databases. Particularly the data concerning pathogenic bacteria, as it can have significant implications for pathogen surveillance, epidemiological investigations, and the identification of antimicrobial resistance in the environment. Therefore, it is essential to thoroughly assess the quality of genomic data related to public health concerns and, if feasible, take steps to enhance its quality before submission. Several studies have been dedicated to improving the quality of low-quality genomes generated through short read sequencing methods [29]. Thus, it is highly recommended to carefully evaluate and, if necessary, improve the quality of genomic data before uploading it to public databases.

## Conclusion

In conclusion, relying on a single quality parameter is not sufficient for accurate genome assessment. Therefore, it is crucial to integrate multiple metrics such as completeness, contiguity, and accuracy to obtain a more comprehensive evaluation of the quality of pathogenic bacterial genomes. Additionally, implementing automated screening procedures for quality checks on uploaded genomic data in public databases is necessary to prevent the dissemination of low-quality sequencing data. Future research should focus on developing a supervised machine learning algorithm that can utilize rankings of quality metrics to classify unknown genomes based on their contiguity, completeness, and accuracy. This approach will facilitate the efficient and precise assessment of genome quality.

---

**Figure S1.**
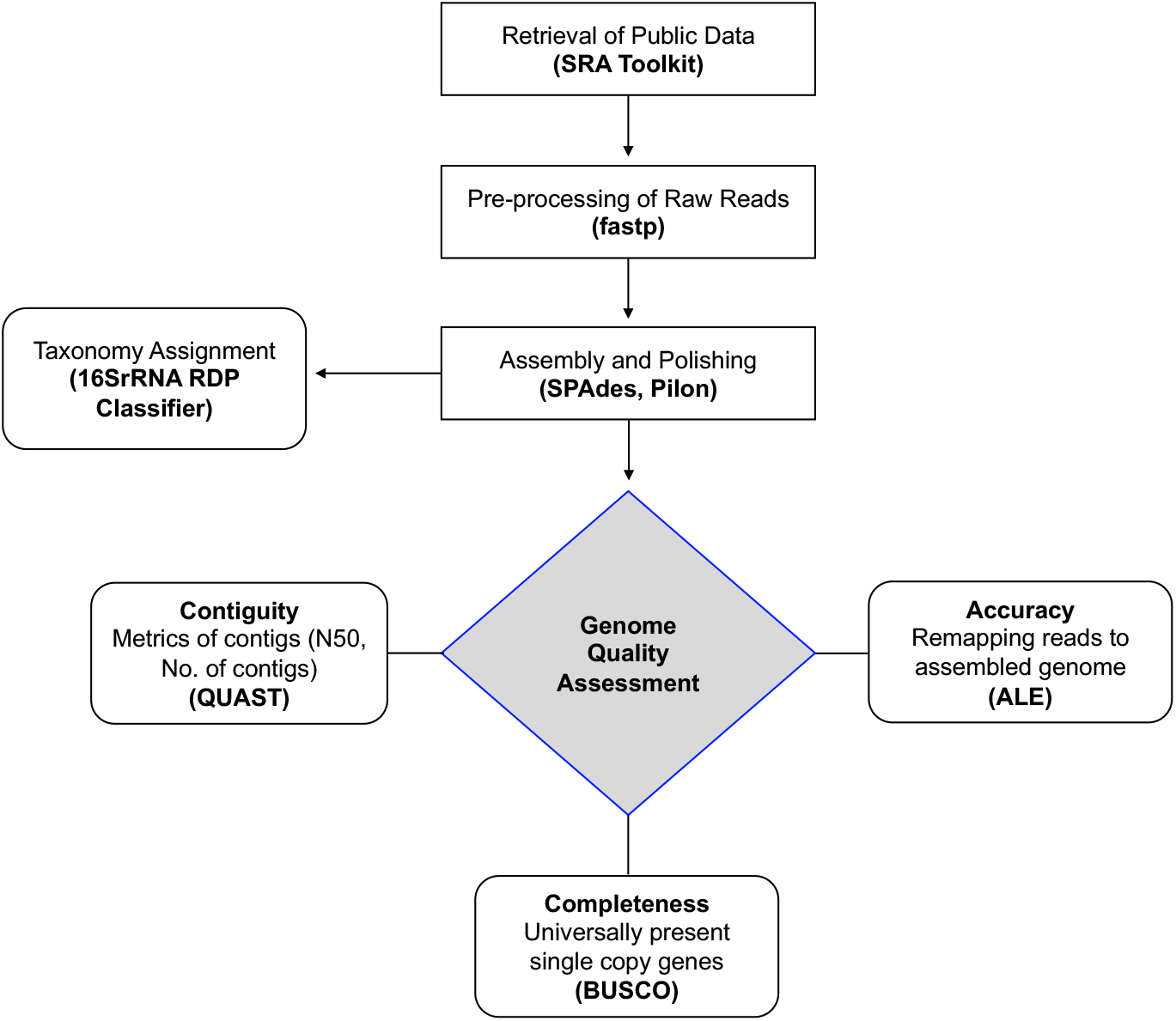
Whole genomes quality assessment workflow; whole genomes downloaded from NCBI pathogen detection database. Low quality reads were screened out using fastp, remaining contigs assembled using SPAdes. Assembled contigs were improved by polishing with reads using Pilon. Taxonomy was determined through 16SRDP classifier. Assembled genomes were assessed for completeness, contiguity, and accuracy by using BUSCO (benchmarking universal single-copy orthologs), QUAST, and ALE (assembly likelihood evaluation), respectively.

