## Supplementary table for "Integrative Assessment of Pathogenic Bacterial Genomes: Insights from Quality Metrics"

**Running Title:** Integrative Assessment of Pathogenic Bacterial Genomes

Table S1. Genomes (n=14) showing contamination of other bacteria predicted through 16S rRNA detection.

| SRA# | Taxonomy (NCBI-PD) | Contamination | Single-copy BUSCO (%) | No. of contigs | N50 | Unmapped Reads |
| --- | --- | --- | --- | --- | --- | --- |
| SRR1013611 | *Mycobacterium tuberculosis KT-0056* | *Bradyrhizobium sp. SK17* | 69.5 | 2371 | 7592 | 9.445998851 |
| SRR1013613 | *Mycobacterium tuberculosis KT-0057* |  | 37.1 | 1858 | 9562 | 6.362663594 |
| SRR1013567 | *Mycobacterium tuberculosis KT-0023* |  | 37.9 | 1251 | 17265 | 2.959541367 |
| SRR1013595 | *Mycobacterium tuberculosis KT-0048* |  | 33.1 | 1761 | 9393 | 2.530354877 |
| SRR1013612 | *Mycobacterium tuberculosis KT-0056* |  | 46 | 2053 | 8182 | 8.431820395 |
| SRR1013583 | *Mycobacterium tuberculosis KT-0039* |  | 87.6 | 1106 | 16284 | 10.75837884 |
| SRR1013634 | *Mycobacterium tuberculosis KT-0064* |  | 28.2 | 934 | 19362 | 1.297955461 |
| SRR1013566 | *Mycobacterium tuberculosis KT-0022* |  | 41.1 | 1967 | 9361 | 7.510776021 |
| SRR1013602 | *Mycobacterium tuberculosis KT-0051* |  | 27.4 | 1105 | 16777 | 1.403999549 |
| SRR1013565 | *Mycobacterium tuberculosis KT-0022* |  | 41.1 | 2297 | 8573 | 8.883631482 |
| SRR1013614 | *Mycobacterium tuberculosis KT-0057* |  | 37.9 | 1867 | 9572 | 6.41192597 |
| SRR1013601 | *Mycobacterium tuberculosis KT-0051* |  | 29.8 | 1062 | 17448 | 1.399125528 |
| SRR1013584 | *Mycobacterium tuberculosis KT-0039* |  | 45.2 | 871 | 17659 | 9.100506846 |
| SRR1013633 | *Mycobacterium tuberculosis KT-0064* |  | 30.6 | 909 | 20258 | 1.054034643 |
