## Supplementary File 1 for "Integrative Assessment of Pathogenic Bacterial Genomes: Insights from Quality Metrics"

SRA (Sequence Read Archive) Accession numbers of the bacterial genomes used in the study.

| **SRA#** | **NCBI Taxonomy** |
| --- | --- |
| SRR1144796 | Mycobacterium tuberculosis KT-0034 |
| SRR1013679 | Mycobacterium tuberculosis KT-0110 |
| SRR11486432 | Listeria monocytogenes |
| SRR1013682 | Mycobacterium tuberculosis KT-0107 |
| SRR1013573 | Mycobacterium tuberculosis KT-0027 |
| SRR11486365 | Listeria monocytogenes |
| SRR6784331 | Listeria monocytogenes |
| SRR7136055 | Acinetobacter baumannii |
| SRR1049000 | Mycobacterium tuberculosis KT-0058 |
| SRR1013572 | Mycobacterium tuberculosis KT-0026 |
| SRR1048995 | Mycobacterium tuberculosis KT-0056 |
| SRR1049028 | Mycobacterium tuberculosis KT-0075 |
| SRR1805599 | Salmonella enterica subsp. enterica |
| SRR5434331 | Escherichia coli |
| SRR1013625 | Mycobacterium tuberculosis KT-0071 |
| SRR7136088 | Acinetobacter baumannii |
| SRR1048970 | Mycobacterium tuberculosis KT-0041 |
| SRR11787127 | Listeria monocytogenes |
| SRR7136048 | Acinetobacter baumannii |
| SRR1013611 | Mycobacterium tuberculosis KT-0056 |
| SRR1048977 | Mycobacterium tuberculosis KT-0045 |
| SRR1013638 | Mycobacterium tuberculosis KT-0077 |
| SRR7235684 | Cronobacter sakazakii |
| SRR11787108 | Listeria monocytogenes |
| SRR1048855 | Mycobacterium tuberculosis KT-0014 |
| SRR1144807 | Mycobacterium tuberculosis KT-0025 |
| SRR7235689 | Cronobacter sakazakii |
| SRR7136113 | Acinetobacter baumannii |
| SRR1049042 | Mycobacterium tuberculosis KT-0083 |
| SRR7136094 | Acinetobacter baumannii |
| SRR7136106 | Acinetobacter baumannii |
| SRR7209071 | Cronobacter sakazakii |
| SRR7136034 | Acinetobacter baumannii |
| SRR11787130 | Listeria monocytogenes |
| SRR1048980 | Mycobacterium tuberculosis KT-0048 |
| SRR9853553 | Salmonella enterica subsp. enterica serovar Agona |
| SRR7359081 | Listeria monocytogenes |
| SRR7136039 | Acinetobacter baumannii |
| SRR2878463 | Salmonella enterica subsp. enterica serovar Newport |
| SRR1048836 | Mycobacterium tuberculosis KT-0001 |
| SRR7136089 | Acinetobacter baumannii |
| SRR1144768 | Mycobacterium tuberculosis KT-0060 |
| SRR1048847 | Mycobacterium tuberculosis KT-0008 |
| SRR1049039 | Mycobacterium tuberculosis KT-0080 |
| SRR7136073 | Acinetobacter baumannii |
| SRR1013639 | Mycobacterium tuberculosis KT-0077 |
| SRR4428965 | Salmonella enterica subsp. enterica serovar Senftenberg |
| SRR1048010 | Mycobacterium tuberculosis KT-0086 |
| SRR11787116 | Listeria monocytogenes |
| SRR1144776 | Mycobacterium tuberculosis KT-0004 |
| SRR11486381 | Listeria monocytogenes |
| SRR1013571 | Mycobacterium tuberculosis KT-0026 |
| SRR1003102 | Mycobacterium tuberculosis SK-C |
| SRR1049034 | Mycobacterium tuberculosis KT-0078 |
| SRR1013667 | Mycobacterium tuberculosis KT-0100 |
| SRR7136099 | Acinetobacter baumannii |
| SRR7235878 | Cronobacter sakazakii |
| SRR1013648 | Mycobacterium tuberculosis KT-0084 |
| SRR1013620 | Mycobacterium tuberculosis KT-0063 |
| SRR7136030 | Acinetobacter baumannii |
| SRR1013642 | Mycobacterium tuberculosis KT-0079 |
| SRR7136112 | Acinetobacter baumannii |
| SRR1013672 | Mycobacterium tuberculosis KT-0106 |
| SRR1013627 | Mycobacterium tuberculosis KT-0072 |
| SRR7136085 | Acinetobacter baumannii |
| SRR1013654 | Mycobacterium tuberculosis KT-0091 |
| SRR1049010 | Mycobacterium tuberculosis KT-0070 |
| SRR2910877 | Salmonella enterica subsp. enterica serovar Senftenberg |
| SRR7136086 | Acinetobacter baumannii |
| SRR1048973 | Mycobacterium tuberculosis KT-0043 |
| SRR1049063 | Mycobacterium tuberculosis KT-0102 |
| SRR11787109 | Listeria monocytogenes |
| SRR1049046 | Mycobacterium tuberculosis KT-0085 |
| SRR1049009 | Mycobacterium tuberculosis KT-0070 |
| SRR7136124 | Acinetobacter baumannii |
| SRR7209078 | Cronobacter sakazakii |
| SRR7547837 | Listeria monocytogenes |
| SRR1048866 | Mycobacterium tuberculosis KT-0022 |
| SRR1013568 | Mycobacterium tuberculosis KT-0023 |
| SRR1013555 | Mycobacterium tuberculosis KT-0014 |
| SRR7136097 | Acinetobacter baumannii |
| SRR5434334 | Escherichia coli |
| SRR3438086 | Salmonella enterica subsp. enterica serovar Oranienburg |
| SRR1013664 | Mycobacterium tuberculosis KT-0099 |
| SRR7136105 | Acinetobacter baumannii |
| SRR1144787 | Mycobacterium tuberculosis KT-0031 |
| SRR1013596 | Mycobacterium tuberculosis KT-0048 |
| SRR7136029 | Acinetobacter baumannii |
| SRR7136044 | Acinetobacter baumannii |
| SRR1049052 | Mycobacterium tuberculosis KT-0092 |
| SRR1048996 | Mycobacterium tuberculosis KT-0056 |
| SRR1048990 | Mycobacterium tuberculosis KT-0053 |
| SRR1823703 | Salmonella enterica subsp. enterica |
| SRR1049062 | Mycobacterium tuberculosis KT-0102 |
| SRR1013585 | Mycobacterium tuberculosis KT-0041 |
| SRR1013543 | Mycobacterium tuberculosis KT-0006 |
| SRR1048857 | Mycobacterium tuberculosis KT-0015 |
| SRR1013626 | Mycobacterium tuberculosis KT-0071 |
| SRR2585813 | Salmonella enterica subsp. enterica serovar Lexington |
| SRR1144749 | Mycobacterium tuberculosis KT-0037 |
| SRR1048864 | Mycobacterium tuberculosis KT-0019 |
| SRR1013645 | Mycobacterium tuberculosis KT-0080 |
| SRR5571307 | Listeria monocytogenes |
| SRR1144813 | Mycobacterium tuberculosis KT-0018 |
| SRR7136120 | Acinetobacter baumannii |
| SRR6926203 | Listeria monocytogenes |
| SRR10887301 | Cronobacter sakazakii |
| SRR7136119 | Acinetobacter baumannii |
| SRR2532675 | Salmonella enterica subsp. enterica serovar Lexington |
| SRR7136043 | Acinetobacter baumannii |
| SRR1048985 | Mycobacterium tuberculosis KT-0051 |
| SRR1013644 | Mycobacterium tuberculosis KT-0080 |
| SRR1013668 | Mycobacterium tuberculosis KT-0102 |
| SRR1013623 | Mycobacterium tuberculosis KT-0070 |
| SRR7136058 | Acinetobacter baumannii |
| SRR7136104 | Acinetobacter baumannii |
| SRR7136114 | Acinetobacter baumannii |
| SRR5811620 | Listeria monocytogenes |
| SRR1013613 | Mycobacterium tuberculosis KT-0057 |
| SRR1049061 | Mycobacterium tuberculosis KT-0100 |
| SRR1048880 | Mycobacterium tuberculosis KT-0027 |
| SRR7136069 | Acinetobacter baumannii |
| SRR11648067 | Listeria monocytogenes |
| SRR7136042 | Acinetobacter baumannii |
| SRR1003101 | Mycobacterium tuberculosis SK-B |
| SRR1013547 | Mycobacterium tuberculosis KT-0008 |
| SRR1013558 | Mycobacterium tuberculosis KT-0015 |
| SRR1013552 | Mycobacterium tuberculosis KT-0011 |
| SRR7136046 | Acinetobacter baumannii |
| SRR1013546 | Mycobacterium tuberculosis KT-0007 |
| SRR3438087 | Salmonella enterica subsp. enterica serovar Oranienburg |
| SRR1288379 | Salmonella enterica subsp. enterica serovar 4,[5],12:i:- |
| SRR1013567 | Mycobacterium tuberculosis KT-0023 |
| SRR1049060 | Mycobacterium tuberculosis KT-0100 |
| SRR1049076 | Mycobacterium tuberculosis KT-0099 |
| SRR1048971 | Mycobacterium tuberculosis KT-0041 |
| SRR11787111 | Listeria monocytogenes |
| SRR7136060 | Acinetobacter baumannii |
| SRR1144780 | Mycobacterium tuberculosis KT-0069 |
| SRR1013641 | Mycobacterium tuberculosis KT-0078 |
| SRR7702518 | Listeria monocytogenes |
| SRR1048851 | Mycobacterium tuberculosis KT-0011 |
| SRR8182744 | Acinetobacter baumannii |
| SRR1049005 | Mycobacterium tuberculosis KT-0064 |
| SRR1049073 | Mycobacterium tuberculosis KT-0109 |
| SRR7136102 | Acinetobacter baumannii |
| SRR9062702 | Vagococcus acidifermentans |
| SRR6303974 | Listeria monocytogenes |
| SRR7136079 | Acinetobacter baumannii |
| SRR7136075 | Acinetobacter baumannii |
| SRR1503321 | Salmonella enterica subsp. enterica serovar Havana |
| SRR7547846 | Listeria monocytogenes |
| SRR7136038 | Acinetobacter baumannii |
| SRR7235666 | Cronobacter sakazakii |
| SRR1652534 | Salmonella enterica |
| SRR7136093 | Acinetobacter baumannii |
| SRR1013592 | Mycobacterium tuberculosis KT-0045 |
| SRR7136096 | Acinetobacter baumannii |
| SRR4429026 | Salmonella enterica subsp. enterica serovar Senftenberg |
| SRR10911064 | Cronobacter sakazakii |
| SRR10911063 | Cronobacter sakazakii |
| SRR1013665 | Mycobacterium tuberculosis KT-0099 |
| SRR11787110 | Listeria monocytogenes |
| SRR5434330 | Escherichia coli |
| SRR10419390 | Listeria monocytogenes |
| SRR11787123 | Listeria monocytogenes |
| SRR1144766 | Mycobacterium tuberculosis KT-0082 |
| SRR1049074 | Mycobacterium tuberculosis KT-0110 |
| SRR1013593 | Mycobacterium tuberculosis KT-0047 |
| SRR11787114 | Listeria monocytogenes |
| SRR1182722 | Salmonella enterica subsp. enterica serovar Oranienburg |
| SRR7136051 | Acinetobacter baumannii |
| SRR7136045 | Acinetobacter baumannii |
| SRR1049056 | Mycobacterium tuberculosis KT-0096 |
| SRR7136062 | Acinetobacter baumannii |
| SRR1610005 | Listeria monocytogenes |
| SRR7136081 | Acinetobacter baumannii |
| SRR7136101 | Acinetobacter baumannii |
| SRR1049032 | Mycobacterium tuberculosis KT-0077 |
| SRR7136084 | Acinetobacter baumannii |
| SRR10911065 | Cronobacter sakazakii |
| SRR7136121 | Acinetobacter baumannii |
| SRR1013563 | Mycobacterium tuberculosis KT-0019 |
| SRR7136059 | Acinetobacter baumannii |
| SRR1049033 | Mycobacterium tuberculosis KT-0077 |
| SRR5182484 | Listeria monocytogenes |
| SRR7136057 | Acinetobacter baumannii |
| SRR7136054 | Acinetobacter baumannii |
| SRR1144805 | Mycobacterium tuberculosis KT-0033 |
| SRR7136122 | Acinetobacter baumannii |
| SRR1013656 | Mycobacterium tuberculosis KT-0092 |
| SRR1013595 | Mycobacterium tuberculosis KT-0048 |
| SRR11787117 | Listeria monocytogenes |
| SRR7136064 | Acinetobacter baumannii |
| SRR7136032 | Acinetobacter baumannii |
| SRR1049071 | Mycobacterium tuberculosis KT-0108 |
| SRR7136052 | Acinetobacter baumannii |
| SRR5071100 | Vibrio parahaemolyticus |
| SRR6806305 | Listeria monocytogenes |
| SRR9062699 | Vagococcus humatus |
| SRR1049050 | Mycobacterium tuberculosis KT-0091 |
| SRR1144737 | Mycobacterium tuberculosis KT-0029 |
| SRR10911067 | Cronobacter sakazakii |
| SRR10911062 | Cronobacter sakazakii |
| SRR7136035 | Acinetobacter baumannii |
| SRR1144733 | Mycobacterium tuberculosis KT-0010 |
| SRR2585418 | Salmonella enterica subsp. enterica serovar Havana |
| SRR6304499 | Listeria monocytogenes |
| SRR10887303 | Cronobacter sakazakii |
| SRR7136033 | Acinetobacter baumannii |
| SRR3453146 | Listeria monocytogenes |
| SRR1918970 | Salmonella enterica |
| SRR7826315 | Salmonella enterica subsp. enterica serovar Kottbus |
| SRR1049049 | Mycobacterium tuberculosis KT-0089 |
| SRR7136065 | Acinetobacter baumannii |
| SRR7136078 | Acinetobacter baumannii |
| SRR11787122 | Listeria monocytogenes |
| SRR1049002 | Mycobacterium tuberculosis KT-0063 |
| SRR1049057 | Mycobacterium tuberculosis KT-0096 |
| SRR7136050 | Acinetobacter baumannii |
| SRR1561272 | Klebsiella pneumoniae |
| SRR1144754 | Mycobacterium tuberculosis KT-0040 |
| SRR1013677 | Mycobacterium tuberculosis KT-0109 |
| SRR7136110 | Acinetobacter baumannii |
| SRR3438088 | Salmonella enterica subsp. enterica serovar Oranienburg |
| SRR1049059 | Mycobacterium tuberculosis KT-0098 |
| SRR1049072 | Mycobacterium tuberculosis KT-0109 |
| SRR11787121 | Listeria monocytogenes |
| SRR7136100 | Acinetobacter baumannii |
| SRR1946930 | Salmonella enterica |
| SRR7136053 | Acinetobacter baumannii |
| SRR1047988 | Mycobacterium tuberculosis KT-0033 |
| SRR1144771 | Mycobacterium tuberculosis KT-0086 |
| SRR11787120 | Listeria monocytogenes |
| SRR1050530 | Mycobacterium tuberculosis KT-0010 |
| SRR1709626 | Listeria monocytogenes |
| SRR1049058 | Mycobacterium tuberculosis KT-0098 |
| SRR7235685 | Cronobacter sakazakii |
| SRR10887300 | Cronobacter sakazakii |
| SRR7136074 | Acinetobacter baumannii |
| SRR1816847 | Salmonella enterica subsp. enterica |
| SRR11192678 | Aeromonas hydrophila |
| SRR1013673 | Mycobacterium tuberculosis KT-0106 |
| SRR11787115 | Listeria monocytogenes |
| SRR1049045 | Mycobacterium tuberculosis KT-0084 |
| SRR1049020 | Mycobacterium tuberculosis KT-0071 |
| SRR1509623 | Salmonella enterica subsp. enterica |
| SRR7136067 | Acinetobacter baumannii |
| SRR7136125 | Acinetobacter baumannii |
| SRR7235686 | Cronobacter sakazakii |
| SRR1049043 | Mycobacterium tuberculosis KT-0083 |
| SRR7136111 | Acinetobacter baumannii |
| SRR7136063 | Acinetobacter baumannii |
| SRR1553800 | Salmonella enterica subsp. enterica serovar Typhimurium |
| SRR7136071 | Acinetobacter baumannii |
| SRR1144820 | Mycobacterium tuberculosis KT-0032 |
| SRR1049077 | Mycobacterium tuberculosis KT-0099 |
| SRR7136108 | Acinetobacter baumannii |
| SRR4429027 | Salmonella enterica |
| SRR1803044 | Salmonella enterica subsp. enterica serovar Senftenberg |
| SRR1049036 | Mycobacterium tuberculosis KT-0079 |
| SRR7136061 | Acinetobacter baumannii |
| SRR1144783 | Mycobacterium tuberculosis KT-0081 |
| SRR11787112 | Listeria monocytogenes |
| SRR7136092 | Acinetobacter baumannii |
| SRR3438067 | Salmonella enterica |
| SRR11362441 | Listeria monocytogenes |
| SRR1049075 | Mycobacterium tuberculosis KT-0110 |
| SRR1363336 | Salmonella enterica |
| SRR1144774 | Mycobacterium tuberculosis KT-0020 |
| SRR2078207 | Salmonella enterica subsp. enterica serovar Senftenberg |
| SRR3457705 | Salmonella enterica subsp. enterica serovar Agona |
| SRR1049051 | Mycobacterium tuberculosis KT-0091 |
| SRR7136103 | Acinetobacter baumannii |
| SRR1049024 | Mycobacterium tuberculosis KT-0072 |
| SRR7136080 | Acinetobacter baumannii |
| SRR1049004 | Mycobacterium tuberculosis KT-0064 |
| SRR1049055 | Mycobacterium tuberculosis KT-0094 |
| SRR7136109 | Acinetobacter baumannii |
| SRR7136040 | Acinetobacter baumannii |
| SRR7136082 | Acinetobacter baumannii |
| SRR7136123 | Acinetobacter baumannii |
| SRR7136098 | Acinetobacter baumannii |
| SRR1048998 | Mycobacterium tuberculosis KT-0057 |
| SRR11787129 | Listeria monocytogenes |
| SRR1049044 | Mycobacterium tuberculosis KT-0084 |
| SRR11648057 | Listeria monocytogenes |
| SRR7136049 | Acinetobacter baumannii |
| SRR1685377 | Salmonella enterica subsp. enterica serovar Enteritidis |
| SRR1049048 | Mycobacterium tuberculosis KT-0089 |
| SRR7136126 | Acinetobacter baumannii |
| SRR1049053 | Mycobacterium tuberculosis KT-0092 |
| SRR7136056 | Acinetobacter baumannii |
| SRR4733931 | Salmonella enterica |
| SRR1049066 | Mycobacterium tuberculosis KT-0106 |
| SRR1049069 | Mycobacterium tuberculosis KT-0107 |
| SRR4733932 | Salmonella enterica |
| SRR1049019 | Mycobacterium tuberculosis KT-0071 |
| SRR7136066 | Acinetobacter baumannii |
| SRR2585419 | Salmonella enterica subsp. enterica |
| SRR7235688 | Cronobacter sakazakii |
| SRR1144790 | Mycobacterium tuberculosis KT-0067 |
| SRR11787125 | Listeria monocytogenes |
| SRR7136036 | Acinetobacter baumannii |
| SRR1144817 | Mycobacterium tuberculosis KT-0042 |
| SRR7136047 | Acinetobacter baumannii |
| SRR2124318 | Salmonella enterica subsp. enterica serovar Lexington |
| SRR11787118 | Listeria monocytogenes |
| SRR2174058 | Salmonella enterica subsp. enterica serovar Senftenberg |
| SRR7136117 | Acinetobacter baumannii |
| SRR6919274 | Escherichia coli O57:H16 |
| SRR1726160 | Salmonella enterica subsp. enterica |
| SRR1636908 | Salmonella enterica subsp. enterica serovar Senftenberg |
| SRR1049065 | Mycobacterium tuberculosis KT-0104 |
| SRR1049070 | Mycobacterium tuberculosis KT-0108 |
| SRR1049025 | Mycobacterium tuberculosis KT-0072 |
| SRR1182726 | Salmonella enterica subsp. enterica serovar Oranienburg |
| SRR10887302 | Cronobacter sakazakii |
| SRR1049054 | Mycobacterium tuberculosis KT-0094 |
| SRR2052522 | Escherichia coli |
| SRR7136076 | Acinetobacter baumannii |
| SRR10887299 | Cronobacter sakazakii |
| SRR7136091 | Acinetobacter baumannii |
| SRR7136037 | Acinetobacter baumannii |
| SRR7136031 | Acinetobacter baumannii |
| SRR1049037 | Mycobacterium tuberculosis KT-0079 |
| SRR7136068 | Acinetobacter baumannii |
| SRR6921434 | Escherichia coli O104 |
| SRR7235687 | Cronobacter sakazakii |
| SRR1049038 | Mycobacterium tuberculosis KT-0080 |
| SRR7136095 | Acinetobacter baumannii |
| SRR11787128 | Listeria monocytogenes |
| SRR1730387 | Salmonella enterica subsp. enterica serovar Senftenberg |
| SRR1049068 | Mycobacterium tuberculosis KT-0107 |
| SRR7136087 | Acinetobacter baumannii |
| SRR11787126 | Listeria monocytogenes |
| SRR7136115 | Acinetobacter baumannii |
| SRR7136118 | Acinetobacter baumannii |
| SRR7136070 | Acinetobacter baumannii |
| SRR1049047 | Mycobacterium tuberculosis KT-0085 |
| SRR1049035 | Mycobacterium tuberculosis KT-0078 |
| SRR1144757 | Mycobacterium tuberculosis KT-0012 |
| SRR7136107 | Acinetobacter baumannii |
| SRR1013541 | Mycobacterium tuberculosis KT-0003 |
| SRR1049064 | Mycobacterium tuberculosis KT-0104 |
| SRR7136077 | Acinetobacter baumannii |
| SRR1805641 | Salmonella enterica subsp. enterica serovar Weltevreden |
| SRR7136041 | Acinetobacter baumannii |
| SRR10911066 | Cronobacter sakazakii |
| SRR7136072 | Acinetobacter baumannii |
| SRR1144725 | Mycobacterium tuberculosis KT-0087 |
| SRR1048962 | Mycobacterium tuberculosis KT-0035 |
| SRR1652540 | Salmonella enterica |
| SRR2174092 | Salmonella enterica subsp. enterica serovar Senftenberg |
| SRR11589263 | Acinetobacter baumannii |
| SRR1049029 | Mycobacterium tuberculosis KT-0075 |
| SRR1049067 | Mycobacterium tuberculosis KT-0106 |
| SRR1013580 | Mycobacterium tuberculosis KT-0035 |
| SRR1013559 | Mycobacterium tuberculosis KT-0016 |
| SRR1048997 | Mycobacterium tuberculosis KT-0057 |
| SRR10419414 | Salmonella enterica |
| SRR1013551 | Mycobacterium tuberculosis KT-0011 |
| SRR1013632 | Mycobacterium tuberculosis KT-0075 |
| SRR1013653 | Mycobacterium tuberculosis KT-0089 |
| SRR1013612 | Mycobacterium tuberculosis KT-0056 |
| SRR1013574 | Mycobacterium tuberculosis KT-0027 |
| SRR1048989 | Mycobacterium tuberculosis KT-0053 |
| SRR1013537 | Mycobacterium tuberculosis KT-0001 |
| SRR1013674 | Mycobacterium tuberculosis KT-0108 |
| SRR1048842 | Mycobacterium tuberculosis KT-0006 |
| SRR1048981 | Mycobacterium tuberculosis KT-0048 |
| SRR1048862 | Mycobacterium tuberculosis KT-0017 |
| SRR1048976 | Mycobacterium tuberculosis KT-0045 |
| SRR1048963 | Mycobacterium tuberculosis KT-0035 |
| SRR1013675 | Mycobacterium tuberculosis KT-0108 |
| SRR1013576 | Mycobacterium tuberculosis KT-0028 |
| SRR1013583 | Mycobacterium tuberculosis KT-0039 |
| SRR1013616 | Mycobacterium tuberculosis KT-0058 |
| SRR10419565 | Salmonella enterica |
| SRR1013663 | Mycobacterium tuberculosis KT-0098 |
| SRR1048854 | Mycobacterium tuberculosis KT-0014 |
| SRR1003103 | Mycobacterium tuberculosis SK-C |
| SRR1013544 | Mycobacterium tuberculosis KT-0006 |
| SRR1013659 | Mycobacterium tuberculosis KT-0094 |
| SRR1013619 | Mycobacterium tuberculosis KT-0063 |
| SRR1048871 | Mycobacterium tuberculosis KT-0026 |
| SRR1013643 | Mycobacterium tuberculosis KT-0079 |
| SRR1013634 | Mycobacterium tuberculosis KT-0064 |
| SRR1003106 | Mycobacterium tuberculosis SK-E |
| SRR1013657 | Mycobacterium tuberculosis KT-0092 |
| SRR1013542 | Mycobacterium tuberculosis KT-0003 |
| SRR1013562 | Mycobacterium tuberculosis KT-0017 |
| SRR1013594 | Mycobacterium tuberculosis KT-0047 |
| SRR1013631 | Mycobacterium tuberculosis KT-0075 |
| SRR1048843 | Mycobacterium tuberculosis KT-0006 |
| SRR1048858 | Mycobacterium tuberculosis KT-0016 |
| SRR1013566 | Mycobacterium tuberculosis KT-0022 |
| SRR1013540 | Mycobacterium tuberculosis KT-0002 |
| SRR1013564 | Mycobacterium tuberculosis KT-0019 |
| SRR1013602 | Mycobacterium tuberculosis KT-0051 |
| SRR1049001 | Mycobacterium tuberculosis KT-0063 |
| SRR1013680 | Mycobacterium tuberculosis KT-0110 |
| SRR1013660 | Mycobacterium tuberculosis KT-0096 |
| SRR1013676 | Mycobacterium tuberculosis KT-0109 |
| SRR1013565 | Mycobacterium tuberculosis KT-0022 |
| SRR1013651 | Mycobacterium tuberculosis KT-0085 |
| SRR1048882 | Mycobacterium tuberculosis KT-0028 |
| SRR1013556 | Mycobacterium tuberculosis KT-0014 |
| SRR1013658 | Mycobacterium tuberculosis KT-0094 |
| SRR1013624 | Mycobacterium tuberculosis KT-0070 |
| SRR1048881 | Mycobacterium tuberculosis KT-0028 |
| SRR1013655 | Mycobacterium tuberculosis KT-0091 |
| SRR1048978 | Mycobacterium tuberculosis KT-0047 |
| SRR1013670 | Mycobacterium tuberculosis KT-0104 |
| SRR1013579 | Mycobacterium tuberculosis KT-0035 |
| SRR1013649 | Mycobacterium tuberculosis KT-0084 |
| SRR1013683 | Mycobacterium tuberculosis KT-0107 |
| SRR1048867 | Mycobacterium tuberculosis KT-0023 |
| SRR1013652 | Mycobacterium tuberculosis KT-0089 |
| SRR1048861 | Mycobacterium tuberculosis KT-0017 |
| SRR1013646 | Mycobacterium tuberculosis KT-0083 |
| SRR1013671 | Mycobacterium tuberculosis KT-0104 |
| SRR1048844 | Mycobacterium tuberculosis KT-0007 |
| SRR1013662 | Mycobacterium tuberculosis KT-0098 |
| SRR1013615 | Mycobacterium tuberculosis KT-0058 |
| SRR1013561 | Mycobacterium tuberculosis KT-0017 |
| SRR1013606 | Mycobacterium tuberculosis KT-0053 |
| SRR1047984 | Mycobacterium tuberculosis KT-0025 |
| SRR1013614 | Mycobacterium tuberculosis KT-0057 |
| SRR1048868 | Mycobacterium tuberculosis KT-0023 |
| SRR1048840 | Mycobacterium tuberculosis KT-0003 |
| SRR1013601 | Mycobacterium tuberculosis KT-0051 |
| SRR1013666 | Mycobacterium tuberculosis KT-0100 |
| SRR1048841 | Mycobacterium tuberculosis KT-0003 |
| SRR1048999 | Mycobacterium tuberculosis KT-0058 |
| SRR1048979 | Mycobacterium tuberculosis KT-0047 |
| SRR1048879 | Mycobacterium tuberculosis KT-0027 |
| SRR1013605 | Mycobacterium tuberculosis KT-0053 |
| SRR1013539 | Mycobacterium tuberculosis KT-0002 |
| SRR1013588 | Mycobacterium tuberculosis KT-0043 |
| SRR1048838 | Mycobacterium tuberculosis KT-0002 |
| SRR1013669 | Mycobacterium tuberculosis KT-0102 |
| SRR10419106 | Listeria monocytogenes |
| SRR1013560 | Mycobacterium tuberculosis KT-0016 |
| SRR1013650 | Mycobacterium tuberculosis KT-0085 |
| SRR1048845 | Mycobacterium tuberculosis KT-0007 |
| SRR1048872 | Mycobacterium tuberculosis KT-0026 |
| SRR1013575 | Mycobacterium tuberculosis KT-0028 |
| SRR1013569 | Mycobacterium tuberculosis KT-0024 |
| SRR1048972 | Mycobacterium tuberculosis KT-0043 |
| SRR1048863 | Mycobacterium tuberculosis KT-0019 |
| SRR1013640 | Mycobacterium tuberculosis KT-0078 |
| SRR1013584 | Mycobacterium tuberculosis KT-0039 |
| SRR1048859 | Mycobacterium tuberculosis KT-0016 |
| SRR1048835 | Mycobacterium tuberculosis KT-0001 |
| SRR1013586 | Mycobacterium tuberculosis KT-0041 |
| SRR1013633 | Mycobacterium tuberculosis KT-0064 |
| SRR1048986 | Mycobacterium tuberculosis KT-0051 |
| SRR1013570 | Mycobacterium tuberculosis KT-0024 |
| SRR1048850 | Mycobacterium tuberculosis KT-0011 |
| SRR1013647 | Mycobacterium tuberculosis KT-0083 |
| SRR1048865 | Mycobacterium tuberculosis KT-0022 |
| SRR1013628 | Mycobacterium tuberculosis KT-0072 |
| SRR1048856 | Mycobacterium tuberculosis KT-0015 |
| SRR1013538 | Mycobacterium tuberculosis KT-0001 |
| SRR1013587 | Mycobacterium tuberculosis KT-0043 |
| SRR1048846 | Mycobacterium tuberculosis KT-0008 |
| SRR1003105 | Mycobacterium tuberculosis SK-E |
| SRR1048839 | Mycobacterium tuberculosis KT-0002 |
| SRR1013661 | Mycobacterium tuberculosis KT-0096 |
| SRR1013557 | Mycobacterium tuberculosis KT-0015 |
| SRR1013591 | Mycobacterium tuberculosis KT-0045 |
| SRR1013545 | Mycobacterium tuberculosis KT-0007 |
| SRR1047983 | Mycobacterium tuberculosis KT-0020 |

**Taxonomy Identification of 474 Genomes**

| **Taxonomy** | **Frequency** |
| --- | --- |
| Mycobacterium tuberculosis | 253 |
| Acinetobacter baumannii | 97 |
| Listeria monocytogenes | 43 |
| Salmonella enterica | 33 |
| Cronobacter sakazakii | 21 |
| Salmonella enterica | 11 |
| Escherichia coli | 6 |
| Mycobacterium tuberculosis | 5 |
| Vagococcus acidifermentans | 1 |
| Vibrio parahaemolyticus | 1 |
| Vagococcus humatus | 1 |
| Klebsiella pneumoniae | 1 |
| Aeromonas hydrophila | 1 |
