## Supplementary File 2 for "Integrative Assessment of Pathogenic Bacterial Genomes: Insights from Quality Metrics"

| **Calculation of Z-scores for each of the attributes.** | | | | | | | | | | | |
| --- | --- | --- | --- | --- | --- | --- | --- | --- | --- | --- | --- |
| **SRA #** | **Single BUSCOs (%)** | **z-score** | **No. of contigs** | **z-score** | **N50** | **z-score** | **Unmapped Reads** | **z-score** | ***No. of contigs** | ***Unmapped reads** | **Summed z-score** |
| SRR1144796 | 99.2 | 0.389823279 | 111 | -0.3689224 | 93723 | -0.363509786 | 0.684431202 | -0.026529334 | 0.368922384 | 0.026529334 | 0.421765211 |
| SRR1013679 | 87.8 | -0.2818726 | 480 | 0.79422374 | 17290 | -0.708226334 | 5.423423059 | 4.712462523 | -0.794223737 | -4.712462523 | -6.496785194 |
| SRR11486432 | 99.3 | 0.395715348 | 58 | -0.5359867 | 109006 | -0.294582713 | 0.988340208 | 0.277379672 | 0.535986732 | -0.277379672 | 0.359739695 |
| SRR1013682 | 83.6 | -0.529339503 | 652 | 1.33639483 | 12344 | -0.730533034 | 6.486614796 | 5.77565426 | -1.336394829 | -5.77565426 | -8.371921626 |
| SRR1013573 | 93.5 | 0.053975339 | 297 | 0.21737891 | 33448 | -0.635352968 | 2.67628267 | 1.965322133 | -0.217378913 | -1.965322133 | -2.764078675 |
| SRR11486365 | 100 | 0.436959832 | 33 | -0.6147907 | 436985 | 1.184618528 | 0.345087348 | -0.365873188 | 0.61479067 | 0.365873188 | 2.602242218 |
| SRR6784331 | 99.7 | 0.419283625 | 36 | -0.6053342 | 475186 | 1.356906896 | 0.634528973 | -0.076431563 | 0.605334197 | 0.076431563 | 2.45795628 |
| SRR7136055 | 99.5 | 0.407499486 | 142 | -0.2712055 | 130442 | -0.197905311 | 2.417192873 | 1.706232337 | 0.271205501 | -1.706232337 | -1.22543266 |
| SRR1049000 | 98.9 | 0.372147072 | 119 | -0.3437051 | 93792 | -0.363198593 | 0.35310731 | -0.357853227 | 0.343705124 | 0.357853227 | 0.710506829 |
| SRR1013572 | 97.6 | 0.295550173 | 304 | 0.23944402 | 34025 | -0.63275067 | 2.208625502 | 1.497664966 | -0.239444015 | -1.497664966 | -2.074309478 |
| SRR1048995 | 30.6 | -3.652136135 | 168 | -0.1892494 | 179837 | 0.024868542 | 1.714610615 | 1.003650078 | 0.189249406 | -1.003650078 | -4.441668266 |
| SRR1049028 | 99.1 | 0.38393121 | 135 | -0.2932706 | 93794 | -0.363189573 | 0.415980009 | -0.294980528 | 0.293270604 | 0.294980528 | 0.608992768 |
| SRR1805599 | 99.1 | 0.38393121 | 65 | -0.5139216 | 272357 | 0.442138241 | 0.788538688 | 0.077578151 | 0.513921629 | -0.077578151 | 1.262412929 |
| SRR5434331 | 99.3 | 0.395715348 | 255 | 0.0849883 | 125454 | -0.220401433 | 1.474145716 | 0.76318518 | -0.084988298 | -0.76318518 | -0.672859562 |
| SRR1013625 | 86.5 | -0.358469499 | 540 | 0.98335319 | 14830 | -0.719321053 | 6.041430458 | 5.330469922 | -0.983353188 | -5.330469922 | -7.391613662 |
| SRR7136088 | 99.4 | 0.401607417 | 60 | -0.5296824 | 160377 | -0.062897005 | 0.291624368 | -0.419336168 | 0.529682417 | 0.419336168 | 1.287728998 |
| SRR1048970 | 99.1 | 0.38393121 | 110 | -0.3720745 | 98633 | -0.341365448 | 0.304442781 | -0.406517755 | 0.372074541 | 0.406517755 | 0.821158059 |
| SRR11787127 | 100 | 0.436959832 | 23 | -0.6463122 | 478483 | 1.371776526 | 0.503213301 | -0.207747235 | 0.646312245 | 0.207747235 | 2.662795838 |
| SRR7136048 | 99.4 | 0.401607417 | 92 | -0.4288134 | 171903 | -0.010914184 | 0.113475248 | -0.597485288 | 0.428813377 | 0.597485288 | 1.416991898 |
| SRR1013611 | 69.5 | -1.360121249 | 2371 | 6.75495359 | 7592 | -0.751964785 | 9.445998851 | 8.735038314 | -6.754953588 | -8.735038314 | -17.60207794 |
| SRR1048977 | 99.1 | 0.38393121 | 109 | -0.3752267 | 110922 | -0.28594146 | 0.282871843 | -0.428088693 | 0.375226699 | 0.428088693 | 0.901305142 |
| SRR1013638 | 95 | 0.142356376 | 323 | 0.29933501 | 28350 | -0.658345196 | 2.4327131 | 1.721752564 | -0.299335008 | -1.721752564 | -2.537076392 |
| SRR7235684 | 98.9 | 0.372147072 | 51 | -0.5580518 | 541493 | 1.65595469 | 0.626290082 | -0.084670455 | 0.558051834 | 0.084670455 | 2.670824051 |
| SRR11787108 | 100 | 0.436959832 | 70 | -0.4981608 | 310485 | 0.614097376 | 0.440779361 | -0.270181175 | 0.498160842 | 0.270181175 | 1.819399225 |
| SRR1048855 | 99.2 | 0.389823279 | 105 | -0.3878353 | 105870 | -0.308726226 | 0.180175239 | -0.530785298 | 0.387835329 | 0.530785298 | 0.99971768 |
| SRR1144807 | 98.8 | 0.366255002 | 118 | -0.3468573 | 103360 | -0.320046448 | 0.670395984 | -0.040564553 | 0.346857281 | 0.040564553 | 0.433630389 |
| SRR7235689 | 98.6 | 0.354470864 | 35 | -0.6084864 | 526435 | 1.588042378 | 0.610240988 | -0.100719548 | 0.608486355 | 0.100719548 | 2.651719145 |
| SRR7136113 | 99.5 | 0.407499486 | 93 | -0.4256612 | 223541 | 0.221975707 | 0.533334992 | -0.177625544 | 0.425661219 | 0.177625544 | 1.232761957 |
| SRR1049042 | 98.8 | 0.366255002 | 78 | -0.4729436 | 130694 | -0.196768778 | 0.981924253 | 0.270963716 | 0.472943582 | -0.270963716 | 0.37146609 |
| SRR7136094 | 99.4 | 0.401607417 | 110 | -0.3720745 | 147913 | -0.119110251 | 0.245411507 | -0.465549029 | 0.372074541 | 0.465549029 | 1.120120737 |
| SRR7136106 | 99.5 | 0.407499486 | 148 | -0.2522926 | 213930 | 0.17862963 | 0.265363403 | -0.445597133 | 0.252292556 | 0.445597133 | 1.284018805 |
| SRR7209071 | 98.6 | 0.354470864 | 27 | -0.6337036 | 2329596 | 9.720386182 | 0.816067084 | 0.105106547 | 0.633703615 | -0.105106547 | 10.60345411 |
| SRR7136034 | 99.5 | 0.407499486 | 125 | -0.3247922 | 143491 | -0.139053686 | 0.268761924 | -0.442198612 | 0.324792179 | 0.442198612 | 1.035436591 |
| SRR11787130 | 100 | 0.436959832 | 24 | -0.6431601 | 510871 | 1.517847981 | 0.447528571 | -0.263431965 | 0.643160087 | 0.263431965 | 2.861399866 |
| SRR1048980 | 29 | -3.746409241 | 226 | -0.0064243 | 200851 | 0.119642704 | 1.007978975 | 0.297018439 | 0.00642427 | -0.297018439 | -3.917360706 |
| SRR9853553 | 89.1 | -0.205275702 | 465 | 0.74694137 | 15647 | -0.715636343 | 1.421924238 | 0.710963701 | -0.746941375 | -0.710963701 | -2.378817121 |
| SRR7359081 | 100 | 0.436959832 | 28 | -0.6305515 | 355456 | 0.816918773 | 0.795603482 | 0.084642946 | 0.630551457 | -0.084642946 | 1.799787116 |
| SRR7136039 | 99.5 | 0.407499486 | 70 | -0.4981608 | 158910 | -0.069513246 | 0.19089147 | -0.520069067 | 0.498160842 | 0.520069067 | 1.356216149 |
| SRR2878463 | 99.3 | 0.395715348 | 74 | -0.4855522 | 377448 | 0.916103762 | 0.652097346 | -0.05886319 | 0.485552212 | 0.05886319 | 1.856234512 |
| SRR1048836 | 99.1 | 0.38393121 | 125 | -0.3247922 | 98631 | -0.341374468 | 0.34159475 | -0.369365786 | 0.324792179 | 0.369365786 | 0.736714707 |
| SRR7136089 | 99.5 | 0.407499486 | 122 | -0.3342487 | 146981 | -0.123313616 | 0.299805007 | -0.41115553 | 0.334248651 | 0.41115553 | 1.029590051 |
| SRR1144768 | 98.9 | 0.372147072 | 117 | -0.3500094 | 98754 | -0.340819732 | 0.624201657 | -0.086758879 | 0.350009439 | 0.086758879 | 0.468095658 |
| SRR1048847 | 99.1 | 0.38393121 | 131 | -0.3058792 | 93792 | -0.363198593 | 0.449963126 | -0.26099741 | 0.305879234 | 0.26099741 | 0.587609261 |
| SRR1049039 | 98 | 0.31911845 | 70 | -0.4981608 | 114080 | -0.271698726 | 2.604133095 | 1.893172559 | 0.498160842 | -1.893172559 | -1.347591994 |
| SRR7136073 | 99.6 | 0.413391555 | 106 | -0.3846832 | 137649 | -0.16540139 | 0.135413592 | -0.575546944 | 0.384683171 | 0.575546944 | 1.208220281 |
| SRR1013639 | 94.6 | 0.1187881 | 370 | 0.44748641 | 28101 | -0.659468198 | 1.867926795 | 1.156966259 | -0.447486411 | -1.156966259 | -2.145132768 |
| SRR4428965 | 98.9 | 0.372147072 | 40 | -0.5927256 | 496187 | 1.451622427 | 0.293001441 | -0.417959095 | 0.592725567 | 0.417959095 | 2.834454161 |
| SRR1048010 | 98.1 | 0.325010519 | 204 | -0.0757717 | 55815 | -0.53447671 | 1.788116826 | 1.077156289 | 0.075771735 | -1.077156289 | -1.210850745 |
| SRR11787116 | 100 | 0.436959832 | 23 | -0.6463122 | 474669 | 1.3545752 | 0.649699417 | -0.061261119 | 0.646312245 | 0.061261119 | 2.499108396 |
| SRR1144776 | 98.9 | 0.372147072 | 109 | -0.3752267 | 103954 | -0.317367479 | 0.860668314 | 0.149707777 | 0.375226699 | -0.149707777 | 0.280298514 |
| SRR11486381 | 100 | 0.436959832 | 34 | -0.6116385 | 343493 | 0.762965061 | 0.485989111 | -0.224971425 | 0.611638512 | 0.224971425 | 2.03653483 |
| SRR1013571 | 97.6 | 0.295550173 | 225 | -0.0095764 | 40967 | -0.601441912 | 2.165400738 | 1.454440202 | 0.009576428 | -1.454440202 | -1.750755513 |
| SRR1003102 | 99.1 | 0.38393121 | 120 | -0.340553 | 101640 | -0.327803731 | 0.462681579 | -0.248278957 | 0.340552966 | 0.248278957 | 0.644959402 |
| SRR1049034 | 99.2 | 0.389823279 | 116 | -0.3531616 | 105431 | -0.310706137 | 0.886027933 | 0.175067397 | 0.353161596 | -0.175067397 | 0.257211342 |
| SRR1013667 | 89.9 | -0.158139149 | 437 | 0.65868096 | 18325 | -0.703558433 | 5.1271274 | 4.416166864 | -0.658680964 | -4.416166864 | -5.93654541 |
| SRR7136099 | 99.4 | 0.401607417 | 71 | -0.4950087 | 160614 | -0.061828123 | 0.151425677 | -0.55953486 | 0.495008684 | 0.55953486 | 1.394322838 |
| SRR7235878 | 98.9 | 0.372147072 | 54 | -0.5485954 | 502355 | 1.479440407 | 1.115293579 | 0.404333043 | 0.548595362 | -0.404333043 | 1.995849797 |
| SRR1013648 | 87.9 | -0.275980531 | 494 | 0.83835394 | 17224 | -0.708523997 | 5.246200016 | 4.53523948 | -0.838353942 | -4.53523948 | -6.35809795 |
| SRR1013620 | 93.9 | 0.077543616 | 393 | 0.51998603 | 22881 | -0.683010652 | 3.680690298 | 2.969729762 | -0.519986034 | -2.969729762 | -4.095182832 |
| SRR7136030 | 99.5 | 0.407499486 | 57 | -0.5391389 | 172043 | -0.010282777 | 0.137924144 | -0.573036392 | 0.539138889 | 0.573036392 | 1.50939199 |
| SRR1013642 | 93.3 | 0.042191201 | 419 | 0.60194213 | 23062 | -0.682194333 | 2.867102373 | 2.156141836 | -0.601942129 | -2.156141836 | -3.398087097 |
| SRR7136112 | 99.5 | 0.407499486 | 94 | -0.4225091 | 169212 | -0.023050725 | 0.329235392 | -0.381725144 | 0.422509062 | 0.381725144 | 1.188682967 |
| SRR1013672 | 81.8 | -0.635396747 | 632 | 1.27335168 | 11655 | -0.733640458 | 7.058498604 | 6.347538067 | -1.273351679 | -6.347538067 | -8.989926951 |
| SRR1013627 | 90 | -0.15224708 | 532 | 0.95813593 | 15269 | -0.717341142 | 4.96027018 | 4.249309644 | -0.958135928 | -4.249309644 | -6.077033793 |
| SRR7136085 | 99.5 | 0.407499486 | 97 | -0.4130526 | 169213 | -0.023046215 | 0.446147838 | -0.264812698 | 0.413052589 | 0.264812698 | 1.062318559 |
| SRR1013654 | 87.2 | -0.317225015 | 504 | 0.86987552 | 17104 | -0.709065203 | 5.773339341 | 5.062378805 | -0.869875517 | -5.062378805 | -6.95854454 |
| SRR1049010 | 99.2 | 0.389823279 | 116 | -0.3531616 | 98756 | -0.340810712 | 0.293639882 | -0.417320655 | 0.353161596 | 0.417320655 | 0.819494819 |
| SRR2910877 | 98.6 | 0.354470864 | 51 | -0.5580518 | 407982 | 1.053813587 | 2.395994837 | 1.685034301 | 0.558051834 | -1.685034301 | 0.281301985 |
| SRR7136086 | 99.5 | 0.407499486 | 99 | -0.4067483 | 169212 | -0.023050725 | 0.345164392 | -0.365796144 | 0.406748274 | 0.365796144 | 1.156993179 |
| SRR1048973 | 98.9 | 0.372147072 | 111 | -0.3689224 | 113484 | -0.274386715 | 0.267216935 | -0.443743602 | 0.368922384 | 0.443743602 | 0.910426342 |
| SRR1049063 | 98.9 | 0.372147072 | 90 | -0.4351177 | 125168 | -0.221691307 | 0.578611265 | -0.132349271 | 0.435117692 | 0.132349271 | 0.717922727 |
| SRR11787109 | 100 | 0.436959832 | 75 | -0.4824001 | 310514 | 0.614228167 | 0.501671643 | -0.209288893 | 0.482400054 | 0.209288893 | 1.742876946 |
| SRR1049046 | 99.1 | 0.38393121 | 126 | -0.32164 | 95837 | -0.353975544 | 0.490684393 | -0.220276143 | 0.321640021 | 0.220276143 | 0.571871831 |
| SRR1049009 | 99.2 | 0.389823279 | 112 | -0.3657702 | 98756 | -0.340810712 | 0.21423984 | -0.496720697 | 0.365770226 | 0.496720697 | 0.91150349 |
| SRR7136124 | 99.5 | 0.407499486 | 54 | -0.5485954 | 270004 | 0.431526097 | 0.158156289 | -0.552804248 | 0.548595362 | 0.552804248 | 1.940425193 |
| SRR7209078 | 98.6 | 0.354470864 | 40 | -0.5927256 | 327867 | 0.692491041 | 0.630112027 | -0.08084851 | 0.592725567 | 0.08084851 | 1.720535982 |
| SRR7547837 | 100 | 0.436959832 | 80 | -0.4666393 | 152586 | -0.098034794 | 1.224520681 | 0.513560144 | 0.466639267 | -0.513560144 | 0.29200416 |
| SRR1048866 | 25 | -3.982092006 | 178 | -0.1577278 | 214680 | 0.182012166 | 0.202089771 | -0.508870765 | 0.157727831 | 0.508870765 | -3.133481243 |
| SRR1013568 | 26.6 | -3.8878189 | 1016 | 2.48378016 | 17996 | -0.705042239 | 1.071034967 | 0.360074431 | -2.483780162 | -0.360074431 | -7.436715732 |
| SRR1013555 | 96 | 0.201277067 | 299 | 0.22368323 | 31268 | -0.645184874 | 1.96911632 | 1.258155784 | -0.223683228 | -1.258155784 | -1.925746818 |
| SRR7136097 | 99.5 | 0.407499486 | 63 | -0.5202259 | 160377 | -0.062897005 | 0.128871596 | -0.582088941 | 0.520225944 | 0.582088941 | 1.446917367 |
| SRR5434334 | 99.3 | 0.395715348 | 279 | 0.16064008 | 116072 | -0.262714709 | 0.691948177 | -0.019012359 | -0.160640078 | 0.019012359 | -0.00862708 |
| SRR3438086 | 98.9 | 0.372147072 | 39 | -0.5958777 | 695628 | 2.351111036 | 1.028842606 | 0.31788207 | 0.595877725 | -0.31788207 | 3.001253763 |
| SRR1013664 | 83.4 | -0.541123642 | 624 | 1.24813442 | 13455 | -0.72552237 | 5.747559278 | 5.036598742 | -1.248134419 | -5.036598742 | -7.551379172 |
| SRR7136105 | 99.5 | 0.407499486 | 92 | -0.4288134 | 223541 | 0.221975707 | 0.124821 | -0.586139536 | 0.428813377 | 0.586139536 | 1.644428106 |
| SRR1144787 | 99.1 | 0.38393121 | 124 | -0.3279443 | 98633 | -0.341365448 | 0.752946835 | 0.041986298 | 0.327944336 | -0.041986298 | 0.3285238 |
| SRR1013596 | 32.3 | -3.55197096 | 1755 | 4.81322456 | 9372 | -0.743936899 | 2.085599688 | 1.374639152 | -4.813224562 | -1.374639152 | -10.48377157 |
| SRR7136029 | 99.4 | 0.401607417 | 97 | -0.4130526 | 107870 | -0.299706128 | 0.226255148 | -0.484705388 | 0.413052589 | 0.484705388 | 0.999659266 |
| SRR7136044 | 99.5 | 0.407499486 | 94 | -0.4225091 | 103444 | -0.319667604 | 0.143700682 | -0.567259854 | 0.422509062 | 0.567259854 | 1.077600798 |
| SRR1049052 | 99.2 | 0.389823279 | 109 | -0.3752267 | 113092 | -0.276154654 | 0.273683662 | -0.437276874 | 0.375226699 | 0.437276874 | 0.926172198 |
| SRR1048996 | 30.6 | -3.652136135 | 165 | -0.1987059 | 180179 | 0.026410978 | 1.736737373 | 1.025776836 | 0.198705878 | -1.025776836 | -4.452796115 |
| SRR1048990 | 99.2 | 0.389823279 | 116 | -0.3531616 | 103944 | -0.317412579 | 1.54774748 | 0.836786943 | 0.353161596 | -0.836786943 | -0.411214647 |
| SRR1823703 | 99.3 | 0.395715348 | 117 | -0.3500094 | 134361 | -0.18023043 | 0.311382039 | -0.399578497 | 0.350009439 | 0.399578497 | 0.965072854 |
| SRR1049062 | 97.7 | 0.301442242 | 78 | -0.4729436 | 136098 | -0.172396476 | 2.548203571 | 1.837243035 | 0.472943582 | -1.837243035 | -1.235253686 |
| SRR1013585 | 87.2 | -0.317225015 | 450 | 0.69965901 | 19790 | -0.696951212 | 5.418995812 | 4.708035276 | -0.699659012 | -4.708035276 | -6.421870515 |
| SRR1013543 | 93.1 | 0.030407063 | 300 | 0.22683539 | 28996 | -0.655431704 | 2.791652376 | 2.08069184 | -0.226835385 | -2.08069184 | -2.932551866 |
| SRR1048857 | 99.2 | 0.389823279 | 112 | -0.3657702 | 123230 | -0.230431781 | 0.289831126 | -0.421129411 | 0.365770226 | 0.421129411 | 0.946291134 |
| SRR1013626 | 87.2 | -0.317225015 | 525 | 0.93607083 | 15632 | -0.715703994 | 6.417545839 | 5.706585303 | -0.936070825 | -5.706585303 | -7.675585137 |
| SRR2585813 | 99.3 | 0.395715348 | 62 | -0.5233781 | 302644 | 0.578734084 | 0.47249306 | -0.238467476 | 0.523378102 | 0.238467476 | 1.73629501 |
| SRR1144749 | 99.2 | 0.389823279 | 132 | -0.3027271 | 83115 | -0.411352382 | 0.733750423 | 0.022789887 | 0.302727076 | -0.022789887 | 0.258408086 |
| SRR1048864 | 98.9 | 0.372147072 | 128 | -0.3153357 | 98633 | -0.341365448 | 0.327201784 | -0.383758752 | 0.315335706 | 0.383758752 | 0.729876083 |
| SRR1013645 | 86.3 | -0.370253637 | 540 | 0.98335319 | 14130 | -0.722478087 | 5.870701328 | 5.159740792 | -0.983353188 | -5.159740792 | -7.235825704 |
| SRR5571307 | 100 | 0.436959832 | 24 | -0.6431601 | 389772 | 0.971685602 | 0.751168801 | 0.040208265 | 0.643160087 | -0.040208265 | 2.011597256 |
| SRR1144813 | 98.9 | 0.372147072 | 122 | -0.3342487 | 95834 | -0.353989074 | 0.773609428 | 0.062648892 | 0.334248651 | -0.062648892 | 0.289757757 |
| SRR7136120 | 99.4 | 0.401607417 | 103 | -0.3941396 | 169212 | -0.023050725 | 0.387562845 | -0.323397691 | 0.394139644 | 0.323397691 | 1.096094027 |
| SRR6926203 | 100 | 0.436959832 | 20 | -0.6557687 | 506431 | 1.497823365 | 0.366629302 | -0.344331235 | 0.655768717 | 0.344331235 | 2.934883149 |
| SRR10887301 | 98.9 | 0.372147072 | 54 | -0.5485954 | 502355 | 1.479440407 | 1.115293579 | 0.404333043 | 0.548595362 | -0.404333043 | 1.995849797 |
| SRR7136119 | 99.5 | 0.407499486 | 74 | -0.4855522 | 199880 | 0.115263446 | 0.213060502 | -0.497900035 | 0.485552212 | 0.497900035 | 1.506215179 |
| SRR2532675 | 99.3 | 0.395715348 | 83 | -0.4571828 | 225994 | 0.233038856 | 0.662103603 | -0.048856934 | 0.457182794 | 0.048856934 | 1.134793933 |
| SRR7136043 | 99.5 | 0.407499486 | 103 | -0.3941396 | 142241 | -0.144691247 | 0.300569308 | -0.410391228 | 0.394139644 | 0.410391228 | 1.067339112 |
| SRR1048985 | 24.2 | -4.029228559 | 159 | -0.2176188 | 183712 | 0.04234498 | 0.433470342 | -0.277490194 | 0.217618823 | 0.277490194 | -3.491774561 |
| SRR1013644 | 87.8 | -0.2818726 | 550 | 1.01487476 | 15047 | -0.718342373 | 6.066086936 | 5.355126399 | -1.014874763 | -5.355126399 | -7.370216135 |
| SRR1013668 | 85.9 | -0.393821914 | 533 | 0.96128809 | 15277 | -0.717305061 | 6.143932393 | 5.432971856 | -0.961288085 | -5.432971856 | -7.505386916 |
| SRR1013623 | 91.9 | -0.040297767 | 370 | 0.44748641 | 22705 | -0.68380442 | 5.002249014 | 4.291288477 | -0.447486411 | -4.291288477 | -5.462877075 |
| SRR7136058 | 99.4 | 0.401607417 | 71 | -0.4950087 | 114711 | -0.268852886 | 0.300415967 | -0.41054457 | 0.495008684 | 0.41054457 | 1.038307785 |
| SRR7136104 | 99.4 | 0.401607417 | 75 | -0.4824001 | 160614 | -0.061828123 | 0.171153037 | -0.539807499 | 0.482400054 | 0.539807499 | 1.361986848 |
| SRR7136114 | 99.5 | 0.407499486 | 89 | -0.4382698 | 147011 | -0.123178315 | 0.623191404 | -0.087769133 | 0.438269849 | 0.087769133 | 0.810360153 |
| SRR5811620 | 100 | 0.436959832 | 25 | -0.6400079 | 389772 | 0.971685602 | 0.780295511 | 0.069334975 | 0.64000793 | -0.069334975 | 1.979318388 |
| SRR1013613 | 37.1 | -3.269151643 | 1858 | 5.13789679 | 9562 | -0.743079989 | 6.362663594 | 5.651703057 | -5.137896786 | -5.651703057 | -14.80183147 |
| SRR1049061 | 99.1 | 0.38393121 | 111 | -0.3689224 | 113484 | -0.274386715 | 0.246209192 | -0.464751345 | 0.368922384 | 0.464751345 | 0.943218223 |
| SRR1048880 | 99.2 | 0.389823279 | 105 | -0.3878353 | 124114 | -0.226444898 | 0.266022822 | -0.444937714 | 0.387835329 | 0.444937714 | 0.996151424 |
| SRR7136069 | 99.4 | 0.401607417 | 69 | -0.501313 | 199104 | 0.111763649 | 0.760516246 | 0.04955571 | 0.501312999 | -0.04955571 | 0.965128355 |
| SRR11648067 | 99.3 | 0.395715348 | 95 | -0.4193569 | 69290 | -0.473703805 | 1.465407903 | 0.754447366 | 0.419356904 | -0.754447366 | -0.413078919 |
| SRR7136042 | 99.5 | 0.407499486 | 77 | -0.4760957 | 142253 | -0.144637126 | 0.148487617 | -0.562472919 | 0.476095739 | 0.562472919 | 1.301431019 |
| SRR1003101 | 99.1 | 0.38393121 | 116 | -0.3531616 | 98633 | -0.341365448 | 0.367386526 | -0.34357401 | 0.353161596 | 0.34357401 | 0.739301369 |
| SRR1013547 | 87.1 | -0.323117084 | 509 | 0.8856363 | 16673 | -0.711009034 | 5.835608713 | 5.124648176 | -0.885636305 | -5.124648176 | -7.044410599 |
| SRR1013558 | 95.7 | 0.18360086 | 326 | 0.30879148 | 30081 | -0.650538302 | 2.207753554 | 1.496793018 | -0.308791481 | -1.496793018 | -2.27252194 |
| SRR1013552 | 93 | 0.024514994 | 420 | 0.60509429 | 21520 | -0.689148828 | 3.533175898 | 2.822215361 | -0.605094287 | -2.822215361 | -4.091943482 |
| SRR7136046 | 99.5 | 0.407499486 | 79 | -0.4697914 | 142253 | -0.144637126 | 0.146279293 | -0.564681243 | 0.469791424 | 0.564681243 | 1.297335028 |
| SRR1013546 | 93.1 | 0.030407063 | 423 | 0.61455076 | 22913 | -0.68286633 | 3.261221778 | 2.550261241 | -0.614550759 | -2.550261241 | -3.817271268 |
| SRR3438087 | 98.9 | 0.372147072 | 44 | -0.5801169 | 354892 | 0.814375105 | 0.624065352 | -0.086895184 | 0.580116937 | 0.086895184 | 1.853534298 |
| SRR1288379 | 98.9 | 0.372147072 | 79 | -0.4697914 | 277750 | 0.466460934 | 0.614895198 | -0.096065339 | 0.469791424 | 0.096065339 | 1.404464768 |
| SRR1013567 | 37.9 | -3.22201509 | 1251 | 3.22453718 | 17265 | -0.708339085 | 2.959541367 | 2.24858083 | -3.224537177 | -2.24858083 | -9.403472182 |
| SRR1049060 | 99.1 | 0.38393121 | 130 | -0.3090314 | 96320 | -0.35179719 | 0.368393012 | -0.342567525 | 0.309031391 | 0.342567525 | 0.683732936 |
| SRR1049076 | 99.1 | 0.38393121 | 73 | -0.4887044 | 158878 | -0.069657568 | 0.93322399 | 0.222263453 | 0.488704369 | -0.222263453 | 0.580714558 |
| SRR1048971 | 99.1 | 0.38393121 | 109 | -0.3752267 | 114843 | -0.268257559 | 0.327652424 | -0.383308113 | 0.375226699 | 0.383308113 | 0.874208462 |
| SRR11787111 | 100 | 0.436959832 | 28 | -0.6305515 | 476576 | 1.363175863 | 0.389316334 | -0.321644202 | 0.630551457 | 0.321644202 | 2.752331354 |
| SRR7136060 | 99.5 | 0.407499486 | 75 | -0.4824001 | 121122 | -0.239938964 | 0.175572571 | -0.535387965 | 0.482400054 | 0.535387965 | 1.185348542 |
| SRR1144780 | 98.8 | 0.366255002 | 121 | -0.3374008 | 114454 | -0.270011968 | 0.976624452 | 0.265663916 | 0.337400809 | -0.265663916 | 0.167979927 |
| SRR1013641 | 95.2 | 0.154140514 | 316 | 0.27726991 | 32168 | -0.64112583 | 2.341420697 | 1.630460161 | -0.277269906 | -1.630460161 | -2.394715382 |
| SRR7702518 | 100 | 0.436959832 | 26 | -0.6368558 | 503454 | 1.48439695 | 0.771054452 | 0.060093916 | 0.636855772 | -0.060093916 | 2.498118639 |
| SRR1048851 | 99.2 | 0.389823279 | 107 | -0.381531 | 105851 | -0.308811917 | 0.275150095 | -0.435810441 | 0.381531014 | 0.435810441 | 0.898352817 |
| SRR8182744 | 99.6 | 0.413391555 | 73 | -0.4887044 | 205369 | 0.140019103 | 0.17956323 | -0.531397306 | 0.488704369 | 0.531397306 | 1.573512334 |
| SRR1049005 | 25.8 | -3.934955453 | 162 | -0.2081624 | 192693 | 0.082849727 | 0.432520154 | -0.278440382 | 0.208162351 | 0.278440382 | -3.365502992 |
| SRR1049073 | 98.9 | 0.372147072 | 120 | -0.340553 | 98633 | -0.341365448 | 0.324242531 | -0.386718006 | 0.340552966 | 0.386718006 | 0.758052596 |
| SRR7136102 | 99.5 | 0.407499486 | 121 | -0.3374008 | 165432 | -0.040098709 | 0.453134858 | -0.257825679 | 0.337400809 | 0.257825679 | 0.962627265 |
| SRR9062702 | 99.3 | 0.395715348 | 63 | -0.5202259 | 154763 | -0.088216418 | 0.90849045 | 0.197529914 | 0.520225944 | -0.197529914 | 0.630194961 |
| SRR6303974 | 99.7 | 0.419283625 | 34 | -0.6116385 | 398162 | 1.00952491 | 0.790870598 | 0.079910062 | 0.611638512 | -0.079910062 | 1.960536984 |
| SRR7136079 | 99.5 | 0.407499486 | 99 | -0.4067483 | 169212 | -0.023050725 | 0.418653208 | -0.292307329 | 0.406748274 | 0.292307329 | 1.083504364 |
| SRR7136075 | 99.5 | 0.407499486 | 60 | -0.5296824 | 181271 | 0.031335951 | 1.071574619 | 0.360614082 | 0.529682417 | -0.360614082 | 0.607903772 |
| SRR1503321 | 98.9 | 0.372147072 | 92 | -0.4288134 | 85361 | -0.401222813 | 0.875755321 | 0.164794785 | 0.428813377 | -0.164794785 | 0.23494285 |
| SRR7547846 | 99.7 | 0.419283625 | 39 | -0.5958777 | 475207 | 1.357001607 | 0.726671791 | 0.015711254 | 0.595877725 | -0.015711254 | 2.356451701 |
| SRR7136038 | 99.5 | 0.407499486 | 100 | -0.4035961 | 166941 | -0.033293046 | 0.214393037 | -0.4965675 | 0.403596116 | 0.4965675 | 1.274370057 |
| SRR7235666 | 98.9 | 0.372147072 | 43 | -0.5832691 | 567916 | 1.775123705 | 0.580109167 | -0.13085137 | 0.583269095 | 0.13085137 | 2.861391241 |
| SRR1652534 | 99.1 | 0.38393121 | 103 | -0.3941396 | 208568 | 0.154446749 | 2.215906738 | 1.504946202 | 0.394139644 | -1.504946202 | -0.572428599 |
| SRR7136093 | 99.4 | 0.401607417 | 65 | -0.5139216 | 160377 | -0.062897005 | 0.121943299 | -0.589017237 | 0.513921629 | 0.589017237 | 1.441649279 |
| SRR1013592 | 91 | -0.093326389 | 395 | 0.52629035 | 19691 | -0.697397707 | 5.044506492 | 4.333545956 | -0.526290349 | -4.333545956 | -5.6505604 |
| SRR7136096 | 99.7 | 0.419283625 | 87 | -0.4445742 | 174537 | 0.000965284 | 0.397366587 | -0.313593949 | 0.444574164 | 0.313593949 | 1.178417022 |
| SRR4429026 | 98.9 | 0.372147072 | 42 | -0.5864213 | 496187 | 1.451622427 | 4.084265799 | 3.373305263 | 0.586421252 | -3.373305263 | -0.963114512 |
| SRR10911064 | 98.9 | 0.372147072 | 53 | -0.5517475 | 493049 | 1.437469894 | 6.799972126 | 6.089011589 | 0.551747519 | -6.089011589 | -3.727647104 |
| SRR10911063 | 98.9 | 0.372147072 | 51 | -0.5580518 | 541493 | 1.65595469 | 0.626290082 | -0.084670455 | 0.558051834 | 0.084670455 | 2.670824051 |
| SRR1013665 | 86.3 | -0.370253637 | 626 | 1.25443873 | 13044 | -0.727376 | 5.666750567 | 4.95579003 | -1.254438734 | -4.95579003 | -7.307858401 |
| SRR11787110 | 100 | 0.436959832 | 26 | -0.6368558 | 555897 | 1.720917431 | 0.558681408 | -0.152279129 | 0.636855772 | 0.152279129 | 2.947012164 |
| SRR5434330 | 99.3 | 0.395715348 | 116 | -0.3531616 | 142764 | -0.142332491 | 1.033316456 | 0.32235592 | 0.353161596 | -0.32235592 | 0.284188533 |
| SRR10419390 | 99.3 | 0.395715348 | 41 | -0.5895734 | 149002 | -0.114198808 | 0.988372412 | 0.277411876 | 0.58957341 | -0.277411876 | 0.593678074 |
| SRR11787123 | 100 | 0.436959832 | 22 | -0.6494644 | 521263 | 1.564716406 | 0.264556741 | -0.446403796 | 0.649464402 | 0.446403796 | 3.097544436 |
| SRR1144766 | 98.8 | 0.366255002 | 135 | -0.2932706 | 93793 | -0.363194083 | 0.914298064 | 0.203337527 | 0.293270604 | -0.203337527 | 0.092993996 |
| SRR1049074 | 98.8 | 0.366255002 | 109 | -0.3752267 | 123878 | -0.22750927 | 0.202717444 | -0.508243092 | 0.375226699 | 0.508243092 | 1.022215524 |
| SRR1013593 | 88.7 | -0.228843978 | 565 | 1.06215713 | 14169 | -0.722302195 | 4.336805915 | 3.625845379 | -1.062157126 | -3.625845379 | -5.639148678 |
| SRR11787114 | 100 | 0.436959832 | 24 | -0.6431601 | 510869 | 1.517838961 | 0.338389837 | -0.372570699 | 0.643160087 | 0.372570699 | 2.970529579 |
| SRR1182722 | 99.1 | 0.38393121 | 44 | -0.5801169 | 521402 | 1.565343303 | 0.661683362 | -0.049277174 | 0.580116937 | 0.049277174 | 2.578668624 |
| SRR7136051 | 99.6 | 0.413391555 | 139 | -0.280662 | 65863 | -0.489159741 | 0.267929696 | -0.44303084 | 0.280661974 | 0.44303084 | 0.647924628 |
| SRR7136045 | 99.5 | 0.407499486 | 78 | -0.4729436 | 142252 | -0.144641636 | 0.146603926 | -0.56435661 | 0.472943582 | 0.56435661 | 1.300158042 |
| SRR1049056 | 99.1 | 0.38393121 | 110 | -0.3720745 | 114756 | -0.268649933 | 0.321755347 | -0.389205189 | 0.372074541 | 0.389205189 | 0.876561007 |
| SRR7136062 | 99.5 | 0.407499486 | 70 | -0.4981608 | 199204 | 0.112214654 | 0.187234412 | -0.523726124 | 0.498160842 | 0.523726124 | 1.541601106 |
| SRR1610005 | 99.7 | 0.419283625 | 32 | -0.6179428 | 330892 | 0.706133938 | 0.508505886 | -0.20245465 | 0.617942827 | 0.20245465 | 1.94581504 |
| SRR7136081 | 99.5 | 0.407499486 | 105 | -0.3878353 | 211163 | 0.166150325 | 0.563078295 | -0.147882242 | 0.387835329 | 0.147882242 | 1.109367382 |
| SRR7136101 | 99.5 | 0.407499486 | 105 | -0.3878353 | 166406 | -0.035705922 | 0.456465412 | -0.254495124 | 0.387835329 | 0.254495124 | 1.014124018 |
| SRR1049032 | 99.1 | 0.38393121 | 142 | -0.2712055 | 93795 | -0.363185063 | 0.367257803 | -0.343702733 | 0.271205501 | 0.343702733 | 0.635654381 |
| SRR7136084 | 99.5 | 0.407499486 | 92 | -0.4288134 | 169212 | -0.023050725 | 0.348189926 | -0.362770611 | 0.428813377 | 0.362770611 | 1.176032748 |
| SRR10911065 | 98.9 | 0.372147072 | 30 | -0.6242471 | 1007998 | 3.759914928 | 0.105915354 | -0.605045182 | 0.624247142 | 0.605045182 | 5.361354324 |
| SRR7136121 | 99.4 | 0.401607417 | 83 | -0.4571828 | 158912 | -0.069504226 | 0.20876756 | -0.502192976 | 0.457182794 | 0.502192976 | 1.291478961 |
| SRR1013563 | 88.2 | -0.258304324 | 528 | 0.9455273 | 15610 | -0.715803215 | 5.990731354 | 5.279770817 | -0.945527298 | -5.279770817 | -7.199405654 |
| SRR7136059 | 99.5 | 0.407499486 | 94 | -0.4225091 | 104075 | -0.316821763 | 0.390995573 | -0.319964964 | 0.422509062 | 0.319964964 | 0.833151749 |
| SRR1049033 | 99.1 | 0.38393121 | 116 | -0.3531616 | 93792 | -0.363198593 | 0.249457141 | -0.461503396 | 0.353161596 | 0.461503396 | 0.835397609 |
| SRR5182484 | 100 | 0.436959832 | 18 | -0.662073 | 634585 | 2.075804138 | 1.276001582 | 0.565041046 | 0.662073032 | -0.565041046 | 2.609795956 |
| SRR7136057 | 99.5 | 0.407499486 | 63 | -0.5202259 | 175211 | 0.004005057 | 0.139557926 | -0.57140261 | 0.520225944 | 0.57140261 | 1.503133097 |
| SRR7136054 | 99.5 | 0.407499486 | 111 | -0.3689224 | 173112 | -0.005461535 | 2.325362789 | 1.614402252 | 0.368922384 | -1.614402252 | -0.843441918 |
| SRR1144805 | 98.9 | 0.372147072 | 133 | -0.2995749 | 98632 | -0.341369958 | 0.721342205 | 0.010381669 | 0.299574919 | -0.010381669 | 0.319970364 |
| SRR7136122 | 99.5 | 0.407499486 | 78 | -0.4729436 | 223539 | 0.221966687 | 0.202164298 | -0.508796238 | 0.472943582 | 0.508796238 | 1.611205993 |
| SRR1013656 | 90.3 | -0.134570872 | 418 | 0.59878997 | 18590 | -0.70236327 | 4.102547073 | 3.391586536 | -0.598789972 | -3.391586536 | -4.827310651 |
| SRR1013595 | 33.1 | -3.504834407 | 1761 | 4.83213751 | 9393 | -0.743842188 | 2.530354877 | 1.81939434 | -4.832137507 | -1.81939434 | -10.90020844 |
| SRR11787117 | 99.7 | 0.419283625 | 26 | -0.6368558 | 475231 | 1.357109848 | 0.719265145 | 0.008304608 | 0.636855772 | -0.008304608 | 2.404944636 |
| SRR7136064 | 99.5 | 0.407499486 | 61 | -0.5265303 | 158600 | -0.070911361 | 0.412820353 | -0.298140183 | 0.526530259 | 0.298140183 | 1.161258568 |
| SRR7136032 | 99.6 | 0.413391555 | 89 | -0.4382698 | 169830 | -0.020263515 | 0.18268663 | -0.528273907 | 0.438269849 | 0.528273907 | 1.359671796 |
| SRR1049071 | 98.9 | 0.372147072 | 128 | -0.3153357 | 98633 | -0.341365448 | 0.406987907 | -0.30397263 | 0.315335706 | 0.30397263 | 0.65008996 |
| SRR7136052 | 99.5 | 0.407499486 | 89 | -0.4382698 | 171903 | -0.010914184 | 0.157382915 | -0.553577622 | 0.438269849 | 0.553577622 | 1.388432773 |
| SRR5071100 | 99.9 | 0.431067763 | 84 | -0.4540306 | 268851 | 0.426326011 | 0.63001972 | -0.080940816 | 0.454030637 | 0.080940816 | 1.392365226 |
| SRR6806305 | 100 | 0.436959832 | 18 | -0.662073 | 424531 | 1.128450382 | 0.515245609 | -0.195714928 | 0.662073032 | 0.195714928 | 2.423198174 |
| SRR9062699 | 99.8 | 0.425175694 | 11 | -0.6841381 | 1144960 | 4.377620209 | 0.445231545 | -0.265728992 | 0.684138135 | 0.265728992 | 5.752663029 |
| SRR1049050 | 99.2 | 0.389823279 | 134 | -0.2964228 | 101662 | -0.32770451 | 0.406933258 | -0.304027279 | 0.296422761 | 0.304027279 | 0.662568809 |
| SRR1144737 | 63.1 | -1.737213672 | 76 | -0.4792479 | 105672 | -0.309619215 | 0.427691053 | -0.283269484 | 0.479247897 | 0.283269484 | -1.284315507 |
| SRR10911067 | 98.6 | 0.354470864 | 40 | -0.5927256 | 327867 | 0.692491041 | 0.630112027 | -0.08084851 | 0.592725567 | 0.08084851 | 1.720535982 |
| SRR10911062 | 98.9 | 0.372147072 | 47 | -0.5706605 | 564275 | 1.758702618 | 0.591155013 | -0.119805523 | 0.570660465 | 0.119805523 | 2.821315678 |
| SRR7136035 | 99.5 | 0.407499486 | 76 | -0.4792479 | 223539 | 0.221966687 | 0.278756496 | -0.43220404 | 0.479247897 | 0.43220404 | 1.54091811 |
| SRR1144733 | 99.1 | 0.38393121 | 124 | -0.3279443 | 93792 | -0.363198593 | 0.665862993 | -0.045097543 | 0.327944336 | 0.045097543 | 0.393774496 |
| SRR2585418 | 98.9 | 0.372147072 | 48 | -0.5675083 | 447134 | 1.230391011 | 1.336430711 | 0.625470175 | 0.567508307 | -0.625470175 | 1.544576215 |
| SRR6304499 | 100 | 0.436959832 | 27 | -0.6337036 | 420959 | 1.112340488 | 0.680181324 | -0.030779212 | 0.633703615 | 0.030779212 | 2.213783147 |
| SRR10887303 | 98.6 | 0.354470864 | 39 | -0.5958777 | 1016092 | 3.796419262 | 0.543653087 | -0.167307449 | 0.595877725 | 0.167307449 | 4.9140753 |
| SRR7136033 | 99.5 | 0.407499486 | 121 | -0.3374008 | 158910 | -0.069513246 | 0.226438566 | -0.48452197 | 0.337400809 | 0.48452197 | 1.159909019 |
| SRR3453146 | 100 | 0.436959832 | 19 | -0.6589209 | 593137 | 1.888871642 | 1.068447367 | 0.357486831 | 0.658920875 | -0.357486831 | 2.627265518 |
| SRR1918970 | 99.1 | 0.38393121 | 95 | -0.4193569 | 219002 | 0.201504596 | 0.966243106 | 0.255282569 | 0.419356904 | -0.255282569 | 0.749510141 |
| SRR7826315 | 99.1 | 0.38393121 | 85 | -0.4508785 | 224833 | 0.22780269 | 0.596879032 | -0.114081504 | 0.450878479 | 0.114081504 | 1.176693883 |
| SRR1049049 | 98.9 | 0.372147072 | 129 | -0.3121835 | 104207 | -0.316226436 | 0.239014054 | -0.471946483 | 0.312183549 | 0.471946483 | 0.840050666 |
| SRR7136065 | 99.5 | 0.407499486 | 94 | -0.4225091 | 162558 | -0.053060589 | 0.290890222 | -0.420070315 | 0.422509062 | 0.420070315 | 1.197018274 |
| SRR7136078 | 99.5 | 0.407499486 | 101 | -0.400444 | 181124 | 0.030672974 | 0.320271832 | -0.390688705 | 0.400443959 | 0.390688705 | 1.229305124 |
| SRR11787122 | 100 | 0.436959832 | 16 | -0.6683773 | 725691 | 2.486696628 | 0.423823547 | -0.287136989 | 0.668377347 | 0.287136989 | 3.879170797 |
| SRR1049002 | 99.1 | 0.38393121 | 113 | -0.3626181 | 96384 | -0.351508547 | 0.194763579 | -0.516196957 | 0.362618069 | 0.516196957 | 0.911237689 |
| SRR1049057 | 98.9 | 0.372147072 | 103 | -0.3941396 | 105673 | -0.309614705 | 0.288811034 | -0.422149502 | 0.394139644 | 0.422149502 | 0.878821513 |
| SRR7136050 | 99.5 | 0.407499486 | 68 | -0.5044652 | 177857 | 0.015938645 | 0.135463063 | -0.575497473 | 0.504465157 | 0.575497473 | 1.503400762 |
| SRR1561272 | 96.1 | 0.207169136 | 295 | 0.2110746 | 34039 | -0.632687529 | 3.457173259 | 2.746212722 | -0.211074598 | -2.746212722 | -3.382805713 |
| SRR1144754 | 99.1 | 0.38393121 | 100 | -0.4035961 | 105645 | -0.309740987 | 0.979876245 | 0.268915709 | 0.403596116 | -0.268915709 | 0.208870631 |
| SRR1013677 | 87.3 | -0.311332946 | 554 | 1.02748339 | 15044 | -0.718355903 | 4.990454307 | 4.279493771 | -1.027483393 | -4.279493771 | -6.336666012 |
| SRR7136110 | 99.5 | 0.407499486 | 113 | -0.3626181 | 169212 | -0.023050725 | 0.815369688 | 0.104409152 | 0.362618069 | -0.104409152 | 0.642657678 |
| SRR3438088 | 98.9 | 0.372147072 | 43 | -0.5832691 | 478669 | 1.372615395 | 2.156783376 | 1.44582284 | 0.583269095 | -1.44582284 | 0.882208721 |
| SRR1049059 | 99.2 | 0.389823279 | 109 | -0.3752267 | 105835 | -0.308884077 | 0.287127493 | -0.423833044 | 0.375226699 | 0.423833044 | 0.879998944 |
| SRR1049072 | 98.9 | 0.372147072 | 113 | -0.3626181 | 101425 | -0.328773392 | 0.278610979 | -0.432349557 | 0.362618069 | 0.432349557 | 0.838341306 |
| SRR11787121 | 100 | 0.436959832 | 24 | -0.6431601 | 510870 | 1.517843471 | 0.46205465 | -0.248905886 | 0.643160087 | 0.248905886 | 2.846869276 |
| SRR7136100 | 99.5 | 0.407499486 | 135 | -0.2932706 | 215131 | 0.184046198 | 0.413338549 | -0.297621987 | 0.293270604 | 0.297621987 | 1.182438275 |
| SRR1946930 | 99.3 | 0.395715348 | 90 | -0.4351177 | 201002 | 0.120323721 | 0.560779061 | -0.150181475 | 0.435117692 | 0.150181475 | 1.101338236 |
| SRR7136053 | 99.5 | 0.407499486 | 87 | -0.4445742 | 169335 | -0.022495989 | 0.339763131 | -0.371197406 | 0.444574164 | 0.371197406 | 1.200775067 |
| SRR1047988 | 98.3 | 0.336794657 | 223 | -0.0158807 | 54204 | -0.541742398 | 1.796298243 | 1.085337706 | 0.015880743 | -1.085337706 | -1.274404705 |
| SRR1144771 | 99.1 | 0.38393121 | 126 | -0.32164 | 103129 | -0.321088269 | 0.947892493 | 0.236931956 | 0.321640021 | -0.236931956 | 0.147551006 |
| SRR11787120 | 100 | 0.436959832 | 44 | -0.5801169 | 264270 | 0.405665478 | 0.831720915 | 0.120760379 | 0.580116937 | -0.120760379 | 1.301981868 |
| SRR1050530 | 98.5 | 0.348578795 | 225 | -0.0095764 | 55673 | -0.535117137 | 2.403425533 | 1.692464997 | 0.009576428 | -1.692464997 | -1.869426911 |
| SRR1709626 | 99.7 | 0.419283625 | 35 | -0.6084864 | 330890 | 0.706124918 | 1.214624546 | 0.503664009 | 0.608486355 | -0.503664009 | 1.230230888 |
| SRR1049058 | 99.2 | 0.389823279 | 136 | -0.2901184 | 92477 | -0.369129307 | 0.47412133 | -0.236839207 | 0.290118446 | 0.236839207 | 0.547651625 |
| SRR7235685 | 98.6 | 0.354470864 | 39 | -0.5958777 | 1016092 | 3.796419262 | 0.543653087 | -0.167307449 | 0.595877725 | 0.167307449 | 4.9140753 |
| SRR10887300 | 98.6 | 0.354470864 | 27 | -0.6337036 | 2329596 | 9.720386182 | 0.816067084 | 0.105106547 | 0.633703615 | -0.105106547 | 10.60345411 |
| SRR7136074 | 99.5 | 0.407499486 | 69 | -0.501313 | 243277 | 0.310986027 | 0.242029334 | -0.468931203 | 0.501312999 | 0.468931203 | 1.688729715 |
| SRR1816847 | 99.3 | 0.395715348 | 109 | -0.3752267 | 160910 | -0.060493149 | 2.796561426 | 2.08560089 | 0.375226699 | -2.08560089 | -1.375151992 |
| SRR11192678 | 98.9 | 0.372147072 | 86 | -0.4477263 | 241114 | 0.301230792 | 0.286678942 | -0.424281594 | 0.447726322 | 0.424281594 | 1.545385779 |
| SRR1013673 | 82.4 | -0.600044333 | 607 | 1.19454774 | 13098 | -0.727132457 | 5.751611946 | 5.04065141 | -1.194547741 | -5.04065141 | -7.562375941 |
| SRR11787115 | 100 | 0.436959832 | 27 | -0.6337036 | 476576 | 1.363175863 | 0.714328779 | 0.003368243 | 0.633703615 | -0.003368243 | 2.430471067 |
| SRR1049045 | 99.1 | 0.38393121 | 116 | -0.3531616 | 103948 | -0.317394539 | 0.315203152 | -0.395757385 | 0.353161596 | 0.395757385 | 0.815455652 |
| SRR1049020 | 99.2 | 0.389823279 | 121 | -0.3374008 | 102140 | -0.325548707 | 0.281717495 | -0.429243041 | 0.337400809 | 0.429243041 | 0.830918422 |
| SRR1509623 | 99.1 | 0.38393121 | 141 | -0.2743577 | 107547 | -0.301162874 | 9.514956435 | 8.803995898 | 0.274357659 | -8.803995898 | -8.446869904 |
| SRR7136067 | 99.5 | 0.407499486 | 62 | -0.5233781 | 180686 | 0.028697573 | 0.953458952 | 0.242498416 | 0.523378102 | -0.242498416 | 0.717076746 |
| SRR7136125 | 99.5 | 0.407499486 | 74 | -0.4855522 | 223539 | 0.221966687 | 0.289990578 | -0.420969958 | 0.485552212 | 0.420969958 | 1.535988343 |
| SRR7235686 | 98.9 | 0.372147072 | 30 | -0.6242471 | 1007998 | 3.759914928 | 0.105915354 | -0.605045182 | 0.624247142 | 0.605045182 | 5.361354324 |
| SRR1049043 | 98.8 | 0.366255002 | 130 | -0.3090314 | 98633 | -0.341365448 | 0.669106586 | -0.041853951 | 0.309031391 | 0.041853951 | 0.375774897 |
| SRR7136111 | 99.5 | 0.407499486 | 96 | -0.4162047 | 223541 | 0.221975707 | 0.382211318 | -0.328749218 | 0.416204747 | 0.328749218 | 1.374429158 |
| SRR7136063 | 99.5 | 0.407499486 | 71 | -0.4950087 | 162307 | -0.054192611 | 0.216028909 | -0.494931628 | 0.495008684 | 0.494931628 | 1.343247187 |
| SRR1553800 | 98.9 | 0.372147072 | 90 | -0.4351177 | 376539 | 0.912004128 | 1.406629338 | 0.695668802 | 0.435117692 | -0.695668802 | 1.023600089 |
| SRR7136071 | 99.5 | 0.407499486 | 63 | -0.5202259 | 169212 | -0.023050725 | 0.819767854 | 0.108807318 | 0.520225944 | -0.108807318 | 0.795867388 |
| SRR1144820 | 98.9 | 0.372147072 | 127 | -0.3184879 | 95833 | -0.353993584 | 0.766484445 | 0.055523908 | 0.318487864 | -0.055523908 | 0.281117443 |
| SRR1049077 | 99.1 | 0.38393121 | 79 | -0.4697914 | 140878 | -0.150838443 | 0.44589638 | -0.265064156 | 0.469791424 | 0.265064156 | 0.967948347 |
| SRR7136108 | 99.5 | 0.407499486 | 141 | -0.2743577 | 215131 | 0.184046198 | 0.353624108 | -0.357336428 | 0.274357659 | 0.357336428 | 1.223239771 |
| SRR4429027 | 99.3 | 0.395715348 | 112 | -0.3657702 | 159349 | -0.067533335 | 0.514126937 | -0.196833599 | 0.365770226 | 0.196833599 | 0.890785839 |
| SRR1803044 | 98.9 | 0.372147072 | 41 | -0.5895734 | 347587 | 0.7814292 | 4.781330567 | 4.070370031 | 0.58957341 | -4.070370031 | -2.327220349 |
| SRR1049036 | 98.9 | 0.372147072 | 122 | -0.3342487 | 98633 | -0.341365448 | 0.250944215 | -0.460016321 | 0.334248651 | 0.460016321 | 0.825046596 |
| SRR7136061 | 99.4 | 0.401607417 | 86 | -0.4477263 | 109957 | -0.290293657 | 0.292755003 | -0.418205534 | 0.447726322 | 0.418205534 | 0.977245616 |
| SRR1144783 | 98.7 | 0.360362933 | 119 | -0.3437051 | 103336 | -0.320154689 | 1.596551369 | 0.885590832 | 0.343705124 | -0.885590832 | -0.501677464 |
| SRR11787112 | 100 | 0.436959832 | 24 | -0.6431601 | 510883 | 1.517902102 | 0.489987128 | -0.220973409 | 0.643160087 | 0.220973409 | 2.81899543 |
| SRR7136092 | 99.5 | 0.407499486 | 85 | -0.4508785 | 223541 | 0.221975707 | 0.694596047 | -0.016364489 | 0.450878479 | 0.016364489 | 1.096718162 |
| SRR3438067 | 98.9 | 0.372147072 | 89 | -0.4382698 | 262572 | 0.398007415 | 0.410496253 | -0.300464284 | 0.438269849 | 0.300464284 | 1.50888862 |
| SRR11362441 | 100 | 0.436959832 | 35 | -0.6084864 | 476633 | 1.363432936 | 0.346199801 | -0.364760735 | 0.608486355 | 0.364760735 | 2.773639858 |
| SRR1049075 | 98.8 | 0.366255002 | 111 | -0.3689224 | 112836 | -0.277309227 | 0.25038537 | -0.460575167 | 0.368922384 | 0.460575167 | 0.918443326 |
| SRR1363336 | 98.9 | 0.372147072 | 85 | -0.4508785 | 262568 | 0.397989375 | 3.816364535 | 3.105403998 | 0.450878479 | -3.105403998 | -1.884389073 |
| SRR1144774 | 99.1 | 0.38393121 | 114 | -0.3594659 | 101677 | -0.32763686 | 0.672503826 | -0.038456711 | 0.359465911 | 0.038456711 | 0.454216972 |
| SRR2078207 | 98.9 | 0.372147072 | 106 | -0.3846832 | 235926 | 0.277832659 | 0.721289167 | 0.01032863 | 0.384683171 | -0.01032863 | 1.024334272 |
| SRR3457705 | 98.6 | 0.354470864 | 53 | -0.5517475 | 257713 | 0.376093089 | 0.88421602 | 0.173255484 | 0.551747519 | -0.173255484 | 1.109055989 |
| SRR1049051 | 99.1 | 0.38393121 | 139 | -0.280662 | 91924 | -0.371623364 | 0.361027405 | -0.349933132 | 0.280661974 | 0.349933132 | 0.642902951 |
| SRR7136103 | 99.4 | 0.401607417 | 64 | -0.5170738 | 160614 | -0.061828123 | 0.130173902 | -0.580786634 | 0.517073787 | 0.580786634 | 1.437639715 |
| SRR1049024 | 98.9 | 0.372147072 | 70 | -0.4981608 | 125168 | -0.221691307 | 1.379153493 | 0.668192957 | 0.498160842 | -0.668192957 | -0.01957635 |
| SRR7136080 | 99.5 | 0.407499486 | 91 | -0.4319655 | 223541 | 0.221975707 | 0.210023892 | -0.500936644 | 0.431965534 | 0.500936644 | 1.562377371 |
| SRR1049004 | 28.2 | -3.793545794 | 137 | -0.2869663 | 192711 | 0.082930908 | 0.366824296 | -0.344136241 | 0.286966289 | 0.344136241 | -3.079512357 |
| SRR1049055 | 99.2 | 0.389823279 | 122 | -0.3342487 | 112836 | -0.277309227 | 0.272500215 | -0.438460321 | 0.334248651 | 0.438460321 | 0.885223025 |
| SRR7136109 | 99.5 | 0.407499486 | 101 | -0.400444 | 223539 | 0.221966687 | 0.51771431 | -0.193246226 | 0.400443959 | 0.193246226 | 1.223156359 |
| SRR7136040 | 99.4 | 0.401607417 | 62 | -0.5233781 | 135729 | -0.174060683 | 0.204885864 | -0.506074672 | 0.523378102 | 0.506074672 | 1.256999508 |
| SRR7136082 | 99.5 | 0.407499486 | 107 | -0.381531 | 169212 | -0.023050725 | 0.520912071 | -0.190048465 | 0.381531014 | 0.190048465 | 0.95602824 |
| SRR7136123 | 99.4 | 0.401607417 | 87 | -0.4445742 | 158627 | -0.07078959 | 0.222536623 | -0.488423913 | 0.444574164 | 0.488423913 | 1.263815904 |
| SRR7136098 | 99.5 | 0.407499486 | 67 | -0.5076173 | 160377 | -0.062897005 | 0.107913821 | -0.603046715 | 0.507617314 | 0.603046715 | 1.455266511 |
| SRR1048998 | 28.2 | -3.793545794 | 128 | -0.3153357 | 192693 | 0.082849727 | 0.945206054 | 0.234245518 | 0.315335706 | -0.234245518 | -3.629605879 |
| SRR11787129 | 100 | 0.436959832 | 20 | -0.6557687 | 508469 | 1.507014844 | 0.350762886 | -0.36019765 | 0.655768717 | 0.36019765 | 2.959941044 |
| SRR1049044 | 99.2 | 0.389823279 | 120 | -0.340553 | 98633 | -0.341365448 | 0.354056092 | -0.356904444 | 0.340552966 | 0.356904444 | 0.745915242 |
| SRR11648057 | 98.7 | 0.360362933 | 73 | -0.4887044 | 93820 | -0.363072312 | 4.526042159 | 3.815081622 | 0.488704369 | -3.815081622 | -3.329086631 |
| SRR7136049 | 99.5 | 0.407499486 | 90 | -0.4351177 | 171903 | -0.010914184 | 0.117876035 | -0.593084501 | 0.435117692 | 0.593084501 | 1.424787495 |
| SRR1685377 | 98.9 | 0.372147072 | 29 | -0.6273993 | 543559 | 1.665272451 | 0.79465197 | 0.083691434 | 0.6273993 | -0.083691434 | 2.581127388 |
| SRR1049048 | 99.1 | 0.38393121 | 134 | -0.2964228 | 93807 | -0.363130942 | 0.294849679 | -0.416110857 | 0.296422761 | 0.416110857 | 0.733333886 |
| SRR7136126 | 99.5 | 0.407499486 | 63 | -0.5202259 | 173222 | -0.00496543 | 0.215237601 | -0.495722935 | 0.520225944 | 0.495722935 | 1.418482936 |
| SRR1049053 | 99.2 | 0.389823279 | 117 | -0.3500094 | 98755 | -0.340815222 | 0.264576625 | -0.446383911 | 0.350009439 | 0.446383911 | 0.845401407 |
| SRR7136056 | 99.6 | 0.413391555 | 100 | -0.4035961 | 92089 | -0.370879206 | 1.299345285 | 0.588384748 | 0.403596116 | -0.588384748 | -0.142276282 |
| SRR4733931 | 98.9 | 0.372147072 | 103 | -0.3941396 | 106522 | -0.305785674 | 0.888844167 | 0.177883631 | 0.394139644 | -0.177883631 | 0.282617411 |
| SRR1049066 | 98.5 | 0.348578795 | 73 | -0.4887044 | 183796 | 0.042723824 | 0.440213782 | -0.270746754 | 0.488704369 | 0.270746754 | 1.150753743 |
| SRR1049069 | 98.7 | 0.360362933 | 122 | -0.3342487 | 103513 | -0.31935641 | 0.271194123 | -0.439766413 | 0.334248651 | 0.439766413 | 0.815021588 |
| SRR4733932 | 98.9 | 0.372147072 | 73 | -0.4887044 | 124329 | -0.225475238 | 1.287502219 | 0.576541683 | 0.488704369 | -0.576541683 | 0.05883452 |
| SRR1049019 | 99.1 | 0.38393121 | 123 | -0.3310965 | 101604 | -0.327966093 | 0.279416026 | -0.43154451 | 0.331096494 | 0.43154451 | 0.818606121 |
| SRR7136066 | 99.4 | 0.401607417 | 66 | -0.5107695 | 147704 | -0.120052851 | 0.144220253 | -0.566740284 | 0.510769472 | 0.566740284 | 1.359064322 |
| SRR2585419 | 98.9 | 0.372147072 | 96 | -0.4162047 | 183575 | 0.041727103 | 1.053963149 | 0.343002613 | 0.416204747 | -0.343002613 | 0.487076309 |
| SRR7235688 | 98.6 | 0.354470864 | 39 | -0.5958777 | 685040 | 2.303358641 | 0.425787751 | -0.285172786 | 0.595877725 | 0.285172786 | 3.538880016 |
| SRR1144790 | 98.8 | 0.366255002 | 115 | -0.3563138 | 103631 | -0.318824225 | 0.817670384 | 0.106709847 | 0.356313754 | -0.106709847 | 0.297034684 |
| SRR11787125 | 100 | 0.436959832 | 27 | -0.6337036 | 504594 | 1.489538406 | 0.340640862 | -0.370319674 | 0.633703615 | 0.370319674 | 2.930521526 |
| SRR7136036 | 99.5 | 0.407499486 | 77 | -0.4760957 | 120925 | -0.240827443 | 0.208166621 | -0.502793915 | 0.476095739 | 0.502793915 | 1.145561697 |
| SRR1144817 | 99.3 | 0.395715348 | 112 | -0.3657702 | 98755 | -0.340815222 | 0.640836428 | -0.070124109 | 0.365770226 | 0.070124109 | 0.490794461 |
| SRR7136047 | 99.4 | 0.401607417 | 72 | -0.4918565 | 146978 | -0.123327146 | 0.246052026 | -0.46490851 | 0.491856527 | 0.46490851 | 1.235045308 |
| SRR2124318 | 99.1 | 0.38393121 | 85 | -0.4508785 | 125355 | -0.220847928 | 1.489563257 | 0.778602721 | 0.450878479 | -0.778602721 | -0.16464096 |
| SRR11787118 | 100 | 0.436959832 | 28 | -0.6305515 | 355455 | 0.816914263 | 0.352707813 | -0.358252724 | 0.630551457 | 0.358252724 | 2.242678275 |
| SRR2174058 | 98.6 | 0.354470864 | 82 | -0.460335 | 170912 | -0.015383642 | 1.929545071 | 1.218584534 | 0.460334952 | -1.218584534 | -0.419162361 |
| SRR7136117 | 99.5 | 0.407499486 | 76 | -0.4792479 | 117264 | -0.257338731 | 0.331089941 | -0.379870595 | 0.479247897 | 0.379870595 | 1.009279247 |
| SRR6919274 | 99.5 | 0.407499486 | 181 | -0.1482714 | 105848 | -0.308825447 | 0.35416021 | -0.356800327 | 0.148271358 | 0.356800327 | 0.603745725 |
| SRR1726160 | 98.9 | 0.372147072 | 61 | -0.5265303 | 322260 | 0.667203198 | 1.103401788 | 0.392441252 | 0.526530259 | -0.392441252 | 1.173439277 |
| SRR1636908 | 99.1 | 0.38393121 | 51 | -0.5580518 | 449178 | 1.239609551 | 0.504751806 | -0.20620873 | 0.558051834 | 0.20620873 | 2.387801325 |
| SRR1049065 | 99.1 | 0.38393121 | 120 | -0.340553 | 98880 | -0.340251466 | 0.266299429 | -0.444661108 | 0.340552966 | 0.444661108 | 0.828893818 |
| SRR1049070 | 98.8 | 0.366255002 | 83 | -0.4571828 | 125210 | -0.221501885 | 0.481472332 | -0.229488205 | 0.457182794 | 0.229488205 | 0.831424116 |
| SRR1049025 | 98.9 | 0.372147072 | 68 | -0.5044652 | 118584 | -0.251385467 | 0.552365876 | -0.158594661 | 0.504465157 | 0.158594661 | 0.783821422 |
| SRR1182726 | 98.9 | 0.372147072 | 58 | -0.5359867 | 218053 | 0.19722456 | 1.215486731 | 0.504526195 | 0.535986732 | -0.504526195 | 0.600832169 |
| SRR10887302 | 98.6 | 0.354470864 | 39 | -0.5958777 | 685040 | 2.303358641 | 0.425787751 | -0.285172786 | 0.595877725 | 0.285172786 | 3.538880016 |
| SRR1049054 | 99.1 | 0.38393121 | 160 | -0.2144667 | 84021 | -0.407266278 | 0.367966667 | -0.342993869 | 0.214466666 | 0.342993869 | 0.534125466 |
| SRR2052522 | 99.5 | 0.407499486 | 117 | -0.3500094 | 115734 | -0.264239106 | 0.209903883 | -0.501056653 | 0.350009439 | 0.501056653 | 0.994326473 |
| SRR7136076 | 99.5 | 0.407499486 | 49 | -0.5643561 | 243276 | 0.310981517 | 0.217027571 | -0.493932965 | 0.564356149 | 0.493932965 | 1.776770118 |
| SRR10887299 | 98.9 | 0.372147072 | 43 | -0.5832691 | 567916 | 1.775123705 | 0.580109167 | -0.13085137 | 0.583269095 | 0.13085137 | 2.861391241 |
| SRR7136091 | 99.4 | 0.401607417 | 102 | -0.3972918 | 144510 | -0.134457946 | 0.196654232 | -0.514306305 | 0.397291801 | 0.514306305 | 1.178747577 |
| SRR7136037 | 99.6 | 0.413391555 | 89 | -0.4382698 | 169830 | -0.020263515 | 0.255545516 | -0.45541502 | 0.438269849 | 0.45541502 | 1.28681291 |
| SRR7136031 | 99.5 | 0.407499486 | 120 | -0.340553 | 157932 | -0.073924074 | 0.464149816 | -0.24681072 | 0.340552966 | 0.24681072 | 0.920939099 |
| SRR1049037 | 98.9 | 0.372147072 | 130 | -0.3090314 | 91862 | -0.371902987 | 0.296847485 | -0.414113052 | 0.309031391 | 0.414113052 | 0.723388528 |
| SRR7136068 | 99.4 | 0.401607417 | 78 | -0.4729436 | 177857 | 0.015938645 | 1.182939569 | 0.471979033 | 0.472943582 | -0.471979033 | 0.418510611 |
| SRR6921434 | 99.3 | 0.395715348 | 263 | 0.11020556 | 83814 | -0.408199858 | 1.748775422 | 1.037814886 | -0.110205558 | -1.037814886 | -1.160504954 |
| SRR7235687 | 98.6 | 0.354470864 | 60 | -0.5296824 | 386264 | 0.955864351 | 3.389887089 | 2.678926552 | 0.529682417 | -2.678926552 | -0.83890892 |
| SRR1049038 | 99.2 | 0.389823279 | 114 | -0.3594659 | 105549 | -0.310173951 | 0.382118777 | -0.328841759 | 0.359465911 | 0.328841759 | 0.767956998 |
| SRR7136095 | 99.5 | 0.407499486 | 55 | -0.5454432 | 243276 | 0.310981517 | 0.544557787 | -0.16640275 | 0.545443204 | 0.16640275 | 1.430326957 |
| SRR11787128 | 100 | 0.436959832 | 22 | -0.6494644 | 478483 | 1.371776526 | 0.409069544 | -0.301890992 | 0.649464402 | 0.301890992 | 2.760091752 |
| SRR1730387 | 99.1 | 0.38393121 | 128 | -0.3153357 | 63317 | -0.500642325 | 8.63118265 | 7.920222113 | 0.315335706 | -7.920222113 | -7.721597523 |
| SRR1049068 | 98.8 | 0.366255002 | 152 | -0.2396839 | 93297 | -0.365431067 | 0.34169645 | -0.369264086 | 0.239683926 | 0.369264086 | 0.609771948 |
| SRR7136087 | 99.5 | 0.407499486 | 92 | -0.4288134 | 132723 | -0.18761789 | 0.506598146 | -0.204362391 | 0.428813377 | 0.204362391 | 0.853057364 |
| SRR11787126 | 100 | 0.436959832 | 22 | -0.6494644 | 478483 | 1.371776526 | 0.45949373 | -0.251466806 | 0.649464402 | 0.251466806 | 2.709667566 |
| SRR7136115 | 99.5 | 0.407499486 | 103 | -0.3941396 | 169212 | -0.023050725 | 0.255400476 | -0.45556006 | 0.394139644 | 0.45556006 | 1.234148466 |
| SRR7136118 | 99.5 | 0.407499486 | 90 | -0.4351177 | 84315 | -0.405940324 | 0.36117718 | -0.349783356 | 0.435117692 | 0.349783356 | 0.78646021 |
| SRR7136070 | 99.5 | 0.407499486 | 100 | -0.4035961 | 171538 | -0.012560352 | 1.161064214 | 0.450103678 | 0.403596116 | -0.450103678 | 0.348431573 |
| SRR1049047 | 99.1 | 0.38393121 | 128 | -0.3153357 | 105529 | -0.310264152 | 0.479740897 | -0.23121964 | 0.315335706 | 0.23121964 | 0.620222403 |
| SRR1049035 | 99.1 | 0.38393121 | 109 | -0.3752267 | 125274 | -0.221213242 | 0.875064639 | 0.164104103 | 0.375226699 | -0.164104103 | 0.373840564 |
| SRR1144757 | 98.8 | 0.366255002 | 125 | -0.3247922 | 95833 | -0.353993584 | 0.696332937 | -0.0146276 | 0.324792179 | 0.0146276 | 0.351681197 |
| SRR7136107 | 99.5 | 0.407499486 | 155 | -0.2302275 | 215131 | 0.184046198 | 0.38167778 | -0.329282756 | 0.230227453 | 0.329282756 | 1.151055894 |
| SRR1013541 | 95.3 | 0.160032583 | 290 | 0.19531381 | 28161 | -0.659197595 | 1.55918093 | 0.848220393 | -0.19531381 | -0.848220393 | -1.542699215 |
| SRR1049064 | 99.1 | 0.38393121 | 119 | -0.3437051 | 101664 | -0.32769549 | 0.25473206 | -0.456228476 | 0.343705124 | 0.456228476 | 0.85616932 |
| SRR7136077 | 99.4 | 0.401607417 | 106 | -0.3846832 | 144510 | -0.134457946 | 0.191594915 | -0.519365622 | 0.384683171 | 0.519365622 | 1.171198264 |
| SRR1805641 | 99.3 | 0.395715348 | 117 | -0.3500094 | 142631 | -0.142932328 | 1.975681023 | 1.264720486 | 0.350009439 | -1.264720486 | -0.661928027 |
| SRR7136041 | 99.4 | 0.401607417 | 63 | -0.5202259 | 135729 | -0.174060683 | 0.171093623 | -0.539866913 | 0.520225944 | 0.539866913 | 1.287639591 |
| SRR10911066 | 98.6 | 0.354470864 | 47 | -0.5706605 | 312287 | 0.622224483 | 2.805679948 | 2.094719412 | 0.570660465 | -2.094719412 | -0.5473636 |
| SRR7136072 | 99.5 | 0.407499486 | 69 | -0.501313 | 177857 | 0.015938645 | 1.761373014 | 1.050412478 | 0.501312999 | -1.050412478 | -0.125661347 |
| SRR1144725 | 99.1 | 0.38393121 | 114 | -0.3594659 | 105557 | -0.310137871 | 0.7120412 | 0.001080664 | 0.359465911 | -0.001080664 | 0.432178587 |
| SRR1048962 | 98.9 | 0.372147072 | 123 | -0.3310965 | 98632 | -0.341369958 | 0.30208053 | -0.408880007 | 0.331096494 | 0.408880007 | 0.770753615 |
| SRR1652540 | 99.3 | 0.395715348 | 63 | -0.5202259 | 236262 | 0.279348036 | 1.477531092 | 0.766570555 | 0.520225944 | -0.766570555 | 0.428718773 |
| SRR2174092 | 98.6 | 0.354470864 | 53 | -0.5517475 | 347401 | 0.780590331 | 1.767578479 | 1.056617943 | 0.551747519 | -1.056617943 | 0.630190772 |
| SRR11589263 | 99.5 | 0.407499486 | 114 | -0.3594659 | 134331 | -0.180365731 | 1.170795307 | 0.45983477 | 0.359465911 | -0.45983477 | 0.126764896 |
| SRR1049029 | 99.1 | 0.38393121 | 125 | -0.3247922 | 93792 | -0.363198593 | 0.36491941 | -0.346041126 | 0.324792179 | 0.346041126 | 0.691565922 |
| SRR1049067 | 98.7 | 0.360362933 | 115 | -0.3563138 | 133049 | -0.186147614 | 0.297957629 | -0.413002907 | 0.356313754 | 0.413002907 | 0.943531981 |
| SRR1013580 | 86.7 | -0.346685361 | 501 | 0.86041904 | 16023 | -0.713940565 | 5.98639596 | 5.275435423 | -0.860419045 | -5.275435423 | -7.196480394 |
| SRR1013559 | 94.6 | 0.1187881 | 331 | 0.32455227 | 25892 | -0.669430895 | 2.317026913 | 1.606066377 | -0.324552268 | -1.606066377 | -2.481261441 |
| SRR1048997 | 30.6 | -3.652136135 | 171 | -0.1797929 | 208372 | 0.153562779 | 0.756787423 | 0.045826886 | 0.179792933 | -0.045826886 | -3.364607309 |
| SRR10419414 | 98.9 | 0.372147072 | 120 | -0.340553 | 113029 | -0.276438787 | 0.461513964 | -0.249446573 | 0.340552966 | 0.249446573 | 0.685707823 |
| SRR1013551 | 92.5 | -0.004945352 | 421 | 0.60824644 | 21935 | -0.687277158 | 3.393483302 | 2.682522766 | -0.608246444 | -2.682522766 | -3.982991719 |
| SRR1013632 | 90.8 | -0.105110527 | 416 | 0.59248566 | 20519 | -0.693663387 | 7.016269185 | 6.305308649 | -0.592485657 | -6.305308649 | -7.696568219 |
| SRR1013653 | 91 | -0.093326389 | 508 | 0.88248415 | 18697 | -0.701880695 | 2.960942387 | 2.249981851 | -0.882484147 | -2.249981851 | -3.927673082 |
| SRR1013612 | 46 | -2.744757491 | 2053 | 5.7525675 | 8182 | -0.749303857 | 8.431820395 | 7.720859858 | -5.7525675 | -7.720859858 | -16.96748871 |
| SRR1013574 | 95.2 | 0.154140514 | 290 | 0.19531381 | 32769 | -0.638415291 | 2.175560019 | 1.464599483 | -0.19531381 | -1.464599483 | -2.14418807 |
| SRR1048989 | 99.2 | 0.389823279 | 92 | -0.4288134 | 103944 | -0.317412579 | 1.781141082 | 1.070180545 | 0.428813377 | -1.070180545 | -0.568956469 |
| SRR1013537 | 90.7 | -0.111002596 | 435 | 0.65237665 | 20576 | -0.693406314 | 5.156858199 | 4.445897662 | -0.652376649 | -4.445897662 | -5.902683221 |
| SRR1013674 | 70.4 | -1.307092627 | 901 | 2.12128205 | 7940 | -0.750395288 | 9.418890107 | 8.707929571 | -2.121282049 | -8.707929571 | -12.88669953 |
| SRR1048842 | 98.9 | 0.372147072 | 107 | -0.381531 | 125274 | -0.221213242 | 0.188305368 | -0.522655169 | 0.381531014 | 0.522655169 | 1.055120012 |
| SRR1048981 | 28.2 | -3.793545794 | 228 | -0.00012 | 178224 | 0.017593833 | 0.833365733 | 0.122405196 | 0.000119955 | -0.122405196 | -3.898237202 |
| SRR1048862 | 98.8 | 0.366255002 | 99 | -0.4067483 | 110778 | -0.286590907 | 7.206592754 | 6.495632218 | 0.406748274 | -6.495632218 | -6.009219848 |
| SRR1048976 | 99.1 | 0.38393121 | 151 | -0.2428361 | 76718 | -0.440203163 | 0.389432763 | -0.321527773 | 0.242836084 | 0.321527773 | 0.508091903 |
| SRR1048963 | 98.9 | 0.372147072 | 135 | -0.2932706 | 95840 | -0.353962013 | 0.321974935 | -0.388985601 | 0.293270604 | 0.388985601 | 0.700441263 |
| SRR1013675 | 70.8 | -1.28352435 | 869 | 2.02041301 | 7564 | -0.752091067 | 10.33111217 | 9.62015163 | -2.020413008 | -9.62015163 | -13.67618006 |
| SRR1013576 | 92.3 | -0.01672949 | 385 | 0.49476877 | 17677 | -0.706480945 | 1.230876434 | 0.519915897 | -0.494768774 | -0.519915897 | -1.737895106 |
| SRR1013583 | 87.6 | -0.293656739 | 1106 | 2.76747434 | 16284 | -0.712763443 | 10.75837884 | 10.0474183 | -2.767474338 | -10.0474183 | -13.82131282 |
| SRR1013616 | 86.7 | -0.346685361 | 551 | 1.01802692 | 15535 | -0.716141469 | 5.207076634 | 4.496116098 | -1.01802692 | -4.496116098 | -6.576969848 |
| SRR10419565 | 98.9 | 0.372147072 | 129 | -0.3121835 | 106733 | -0.304834054 | 0.654411273 | -0.056549263 | 0.312183549 | 0.056549263 | 0.43604583 |
| SRR1013663 | 84.7 | -0.464526743 | 671 | 1.39628582 | 11108 | -0.736107454 | 6.391540435 | 5.680579898 | -1.396285822 | -5.680579898 | -8.277499917 |
| SRR1048854 | 99.2 | 0.389823279 | 110 | -0.3720745 | 125308 | -0.2210599 | 0.279065742 | -0.431894794 | 0.372074541 | 0.431894794 | 0.972732714 |
| SRR1003103 | 80.9 | -0.688425369 | 850 | 1.96052202 | 10610 | -0.738353458 | 7.826780297 | 7.11581976 | -1.960522016 | -7.11581976 | -10.5031206 |
| SRR1013544 | 93.9 | 0.077543616 | 303 | 0.23629186 | 29235 | -0.654353803 | 3.543742609 | 2.832782072 | -0.236291858 | -2.832782072 | -3.645884117 |
| SRR1013659 | 83.4 | -0.541123642 | 680 | 1.42465524 | 12263 | -0.730898348 | 5.159257254 | 4.448296718 | -1.424655239 | -4.448296718 | -7.144973946 |
| SRR1013619 | 92.9 | 0.018622925 | 361 | 0.41911699 | 24519 | -0.675623192 | 3.315111686 | 2.60415115 | -0.419116994 | -2.60415115 | -3.680268411 |
| SRR1048871 | 99.1 | 0.38393121 | 102 | -0.3972918 | 125274 | -0.221213242 | 1.174053481 | 0.463092944 | 0.397291801 | -0.463092944 | 0.096916825 |
| SRR1013643 | 93 | 0.024514994 | 446 | 0.68705038 | 22163 | -0.686248867 | 2.986610016 | 2.275649479 | -0.687050382 | -2.275649479 | -3.624433734 |
| SRR1013634 | 28.2 | -3.793545794 | 934 | 2.22530325 | 19362 | -0.698881513 | 1.297955461 | 0.586994925 | -2.225303247 | -0.586994925 | -7.304725478 |
| SRR1003106 | 99.1 | 0.38393121 | 109 | -0.3752267 | 105669 | -0.309632745 | 0.273010445 | -0.437950091 | 0.375226699 | 0.437950091 | 0.887475254 |
| SRR1013657 | 90.2 | -0.140462942 | 401 | 0.54520329 | 22351 | -0.685400977 | 3.848334117 | 3.13737358 | -0.545203294 | -3.13737358 | -4.508440793 |
| SRR1013542 | 93.7 | 0.065759478 | 358 | 0.40966052 | 24590 | -0.675302979 | 2.822350866 | 2.11139033 | -0.409660521 | -2.11139033 | -3.130594352 |
| SRR1013562 | 93.1 | 0.030407063 | 369 | 0.44433425 | 22853 | -0.683136933 | 7.759526991 | 7.048566455 | -0.444334254 | -7.048566455 | -8.145630578 |
| SRR1013594 | 89.6 | -0.175815356 | 543 | 0.99280966 | 15482 | -0.716380502 | 3.628537107 | 2.917576571 | -0.99280966 | -2.917576571 | -4.802582089 |
| SRR1013631 | 90.8 | -0.105110527 | 382 | 0.4853123 | 22990 | -0.682519056 | 6.697297117 | 5.986336581 | -0.485312301 | -5.986336581 | -7.259278466 |
| SRR1048843 | 98.9 | 0.372147072 | 103 | -0.3941396 | 131720 | -0.192141468 | 0.194415807 | -0.516544729 | 0.394139644 | 0.516544729 | 1.090689976 |
| SRR1048858 | 98.9 | 0.372147072 | 110 | -0.3720745 | 112835 | -0.277313737 | 0.243127826 | -0.46783271 | 0.372074541 | 0.46783271 | 0.934740587 |
| SRR1013566 | 41.1 | -3.033468878 | 1967 | 5.48148195 | 9361 | -0.743986509 | 7.510776021 | 6.799815485 | -5.481481954 | -6.799815485 | -16.05875283 |
| SRR1013540 | 93.8 | 0.071651547 | 316 | 0.27726991 | 27291 | -0.663121337 | 3.849534021 | 3.138573485 | -0.277269906 | -3.138573485 | -4.007313181 |
| SRR1013564 | 87.5 | -0.299548808 | 518 | 0.91400572 | 15543 | -0.716105389 | 5.026469318 | 4.315508782 | -0.914005723 | -4.315508782 | -6.245168701 |
| SRR1013602 | 27.4 | -3.840682347 | 1105 | 2.76432218 | 16777 | -0.710539989 | 1.403999549 | 0.693039013 | -2.764322181 | -0.693039013 | -8.008583529 |
| SRR1049001 | 99.1 | 0.38393121 | 130 | -0.3090314 | 89258 | -0.383647154 | 0.359657486 | -0.35130305 | 0.309031391 | 0.35130305 | 0.660618498 |
| SRR1013680 | 88.6 | -0.234736047 | 459 | 0.72802843 | 21791 | -0.687926605 | 4.841054438 | 4.130093901 | -0.728028429 | -4.130093901 | -5.780784983 |
| SRR1013660 | 84.5 | -0.476310881 | 592 | 1.14726538 | 13987 | -0.723123024 | 6.262693278 | 5.551732742 | -1.147265378 | -5.551732742 | -7.898432025 |
| SRR1013676 | 86.1 | -0.382037775 | 583 | 1.11889596 | 15005 | -0.718531795 | 5.620588267 | 4.90962773 | -1.118895961 | -4.90962773 | -7.129093261 |
| SRR1013565 | 41.1 | -3.033468878 | 2297 | 6.52169393 | 8573 | -0.747540428 | 8.883631482 | 8.172670945 | -6.521693932 | -8.172670945 | -18.47537418 |
| SRR1013651 | 88.3 | -0.252412255 | 445 | 0.68389822 | 17070 | -0.709218544 | 4.499436409 | 3.788475872 | -0.683898224 | -3.788475872 | -5.434004896 |
| SRR1048882 | 98.9 | 0.372147072 | 99 | -0.4067483 | 114760 | -0.268631893 | 0.191247167 | -0.519713369 | 0.406748274 | 0.519713369 | 1.029976821 |
| SRR1013556 | 96.6 | 0.236629482 | 316 | 0.27726991 | 36414 | -0.621976164 | 2.15387212 | 1.442911584 | -0.277269906 | -1.442911584 | -2.105528171 |
| SRR1013658 | 84 | -0.505771227 | 680 | 1.42465524 | 11996 | -0.732102531 | 5.120960701 | 4.410000165 | -1.424655239 | -4.410000165 | -7.072529162 |
| SRR1013624 | 93.5 | 0.053975339 | 397 | 0.53259466 | 22177 | -0.686185726 | 3.079016964 | 2.368056428 | -0.532594664 | -2.368056428 | -3.532861478 |
| SRR1048881 | 98.9 | 0.372147072 | 103 | -0.3941396 | 105417 | -0.310769278 | 0.256864879 | -0.454095657 | 0.394139644 | 0.454095657 | 0.909613095 |
| SRR1013655 | 90.4 | -0.128678803 | 509 | 0.8856363 | 17164 | -0.7087946 | 4.981006289 | 4.270045752 | -0.885636305 | -4.270045752 | -5.99315546 |
| SRR1048978 | 99.2 | 0.389823279 | 110 | -0.3720745 | 98972 | -0.339836541 | 0.200323372 | -0.510637164 | 0.372074541 | 0.510637164 | 0.932698443 |
| SRR1013670 | 91.7 | -0.052081905 | 434 | 0.64922449 | 16192 | -0.713178367 | 1.36455901 | 0.653598473 | -0.649224492 | -0.653598473 | -2.068083237 |
| SRR1013579 | 86 | -0.387929844 | 560 | 1.04639634 | 15085 | -0.718170991 | 5.428094208 | 4.717133672 | -1.046396338 | -4.717133672 | -6.869630845 |
| SRR1013649 | 88.4 | -0.246520186 | 463 | 0.74063706 | 17587 | -0.706886849 | 4.798625027 | 4.08766449 | -0.74063706 | -4.08766449 | -5.781708585 |
| SRR1013683 | 85.7 | -0.405606052 | 649 | 1.32693836 | 12768 | -0.728620773 | 5.356364696 | 4.64540416 | -1.326938356 | -4.64540416 | -7.106569341 |
| SRR1048867 | 25.8 | -3.934955453 | 127 | -0.3184879 | 202212 | 0.12578088 | 0.50343017 | -0.207530366 | 0.318487864 | 0.207530366 | -3.283156343 |
| SRR1013652 | 90.6 | -0.116894665 | 499 | 0.85411473 | 18431 | -0.703080368 | 3.724616581 | 3.013656045 | -0.85411473 | -3.013656045 | -4.687745808 |
| SRR1048861 | 99.2 | 0.389823279 | 120 | -0.340553 | 94976 | -0.357858695 | 5.555758537 | 4.844798 | 0.340552966 | -4.844798 | -4.47228045 |
| SRR1013646 | 78.1 | -0.853403305 | 787 | 1.76193609 | 10136 | -0.740491222 | 7.65257115 | 6.941610614 | -1.761936093 | -6.941610614 | -10.29744123 |
| SRR1013671 | 86.4 | -0.364361568 | 554 | 1.02748339 | 14585 | -0.720426015 | 5.835473909 | 5.124513373 | -1.027483393 | -5.124513373 | -7.236784349 |
| SRR1048844 | 99.1 | 0.38393121 | 134 | -0.2964228 | 93303 | -0.365404007 | 0.325441701 | -0.385518836 | 0.296422761 | 0.385518836 | 0.7004688 |
| SRR1013662 | 84.3 | -0.488095019 | 746 | 1.63269763 | 10993 | -0.73662611 | 5.091817459 | 4.380856922 | -1.632697635 | -4.380856922 | -7.238275687 |
| SRR1013615 | 84.8 | -0.458634674 | 562 | 1.05270065 | 15088 | -0.718157461 | 5.521526142 | 4.810565605 | -1.052700653 | -4.810565605 | -7.040058393 |
| SRR1013561 | 91.3 | -0.075650181 | 398 | 0.53574682 | 21701 | -0.688332509 | 7.551310489 | 6.840349952 | -0.535746821 | -6.840349952 | -8.140079464 |
| SRR1013606 | 95 | 0.142356376 | 349 | 0.3812911 | 32978 | -0.637472691 | 2.962518999 | 2.251558463 | -0.381291103 | -2.251558463 | -3.127965881 |
| SRR1047984 | 94.3 | 0.101111892 | 303 | 0.23629186 | 29178 | -0.654610875 | 3.295288459 | 2.584327922 | -0.236291858 | -2.584327922 | -3.374118763 |
| SRR1013614 | 37.9 | -3.22201509 | 1867 | 5.1662662 | 9572 | -0.743034889 | 6.41192597 | 5.700965433 | -5.166266203 | -5.700965433 | -14.83228161 |
| SRR1048868 | 25.8 | -3.934955453 | 114 | -0.3594659 | 192734 | 0.083034639 | 0.662120494 | -0.048840043 | 0.359465911 | 0.048840043 | -3.44361486 |
| SRR1048840 | 99.1 | 0.38393121 | 129 | -0.3121835 | 91942 | -0.371542183 | 0.373045831 | -0.337914705 | 0.312183549 | 0.337914705 | 0.662487281 |
| SRR1013601 | 29.8 | -3.699272688 | 1062 | 2.62877941 | 17448 | -0.707513746 | 1.399125528 | 0.688164992 | -2.628779408 | -0.688164992 | -7.723730834 |
| SRR1013666 | 90.3 | -0.134570872 | 418 | 0.59878997 | 20015 | -0.695936451 | 5.037963871 | 4.327003334 | -0.598789972 | -4.327003334 | -5.756300629 |
| SRR1048841 | 99.1 | 0.38393121 | 113 | -0.3626181 | 105614 | -0.309880798 | 0.344652633 | -0.366307903 | 0.362618069 | 0.366307903 | 0.802976384 |
| SRR1048999 | 99.1 | 0.38393121 | 123 | -0.3310965 | 98764 | -0.340774631 | 0.338257735 | -0.372702802 | 0.331096494 | 0.372702802 | 0.746955874 |
| SRR1048979 | 99.2 | 0.389823279 | 107 | -0.381531 | 102167 | -0.325426936 | 0.232786991 | -0.478173545 | 0.381531014 | 0.478173545 | 0.924100903 |
| SRR1048879 | 99.1 | 0.38393121 | 123 | -0.3310965 | 98690 | -0.341108375 | 0.342657859 | -0.368302677 | 0.331096494 | 0.368302677 | 0.742222006 |
| SRR1013605 | 94.6 | 0.1187881 | 356 | 0.40335621 | 30948 | -0.646628089 | 3.668515167 | 2.95755463 | -0.403356206 | -2.95755463 | -3.888750826 |
| SRR1013539 | 96 | 0.201277067 | 305 | 0.24259617 | 30687 | -0.647805212 | 3.269050152 | 2.558089616 | -0.242596173 | -2.558089616 | -3.247213933 |
| SRR1013588 | 90.4 | -0.128678803 | 500 | 0.85726689 | 18022 | -0.704924978 | 4.433125178 | 3.722164642 | -0.857266887 | -3.722164642 | -5.41303531 |
| SRR1048838 | 99.1 | 0.38393121 | 131 | -0.3058792 | 95851 | -0.353912403 | 0.364839593 | -0.346120943 | 0.305879234 | 0.346120943 | 0.682018984 |
| SRR1013669 | 84.8 | -0.458634674 | 573 | 1.08737439 | 13099 | -0.727127947 | 6.648277988 | 5.937317451 | -1.087374386 | -5.937317451 | -8.210454458 |
| SRR10419106 | 100 | 0.436959832 | 87 | -0.4445742 | 58311 | -0.523219629 | 1.137064023 | 0.426103487 | 0.444574164 | -0.426103487 | -0.06778912 |
| SRR1013560 | 92.7 | 0.006838786 | 360 | 0.41596484 | 25962 | -0.669115192 | 3.662932964 | 2.951972428 | -0.415964836 | -2.951972428 | -4.030213669 |
| SRR1013650 | 89.1 | -0.205275702 | 477 | 0.78476726 | 15640 | -0.715667914 | 4.233122132 | 3.522161596 | -0.784767265 | -3.522161596 | -5.227872476 |
| SRR1048845 | 99.1 | 0.38393121 | 114 | -0.3594659 | 100824 | -0.331483931 | 0.264942956 | -0.44601758 | 0.359465911 | 0.44601758 | 0.85793077 |
| SRR1048872 | 99.1 | 0.38393121 | 106 | -0.3846832 | 124062 | -0.226679421 | 1.212142764 | 0.501182228 | 0.384683171 | -0.501182228 | 0.040752733 |
| SRR1013575 | 92.5 | -0.004945352 | 371 | 0.45063857 | 18532 | -0.702624853 | 1.524577545 | 0.813617009 | -0.450638569 | -0.813617009 | -1.971825783 |
| SRR1013569 | 27.4 | -3.840682347 | 804 | 1.81552277 | 24876 | -0.674013105 | 1.411795796 | 0.70083526 | -1.81552277 | -0.70083526 | -7.031053482 |
| SRR1048972 | 99.1 | 0.38393121 | 127 | -0.3184879 | 96845 | -0.349429415 | 0.348635039 | -0.362325498 | 0.318487864 | 0.362325498 | 0.715315157 |
| SRR1048863 | 98.9 | 0.372147072 | 115 | -0.3563138 | 112836 | -0.277309227 | 0.31408574 | -0.396874797 | 0.356313754 | 0.396874797 | 0.848026395 |
| SRR1013640 | 95.6 | 0.177708791 | 313 | 0.26781343 | 33405 | -0.6355469 | 2.178467715 | 1.467507179 | -0.267813433 | -1.467507179 | -2.193158721 |
| SRR1013584 | 45.2 | -2.791894044 | 871 | 2.02671732 | 17659 | -0.706562126 | 9.100506846 | 8.389546309 | -2.026717323 | -8.389546309 | -13.9147198 |
| SRR1048859 | 98.8 | 0.366255002 | 120 | -0.340553 | 95838 | -0.353971034 | 0.258094526 | -0.45286601 | 0.340552966 | 0.45286601 | 0.805702946 |
| SRR1048835 | 99.1 | 0.38393121 | 117 | -0.3500094 | 95053 | -0.357511422 | 0.370291332 | -0.340669205 | 0.350009439 | 0.340669205 | 0.717098432 |
| SRR1013586 | 90.7 | -0.111002596 | 425 | 0.62085507 | 18639 | -0.702142278 | 4.689858596 | 3.97889806 | -0.620855074 | -3.97889806 | -5.412898008 |
| SRR1013633 | 30.6 | -3.652136135 | 909 | 2.14649931 | 20258 | -0.694840509 | 1.054034643 | 0.343074107 | -2.146499309 | -0.343074107 | -6.83655006 |
| SRR1048986 | 25 | -3.982092006 | 139 | -0.280662 | 179869 | 0.025012863 | 0.47225341 | -0.238707126 | 0.280661974 | 0.238707126 | -3.437710043 |
| SRR1013570 | 28.2 | -3.793545794 | 805 | 1.81867493 | 23664 | -0.679479284 | 1.394173919 | 0.683213382 | -1.818674928 | -0.683213382 | -6.974913388 |
| SRR1048850 | 99.1 | 0.38393121 | 113 | -0.3626181 | 103935 | -0.31745317 | 0.263051538 | -0.447908999 | 0.362618069 | 0.447908999 | 0.877005108 |
| SRR1013647 | 78.3 | -0.841619166 | 790 | 1.77139257 | 10040 | -0.740924186 | 8.120940453 | 7.409979916 | -1.771392565 | -7.409979916 | -10.76391583 |
| SRR1048865 | 25 | -3.982092006 | 173 | -0.1734886 | 219606 | 0.204228666 | 0.208377861 | -0.502582676 | 0.173488618 | 0.502582676 | -3.101792046 |
| SRR1013628 | 86.9 | -0.334901222 | 529 | 0.94867946 | 15412 | -0.716696205 | 5.220247848 | 4.509287311 | -0.948679455 | -4.509287311 | -6.509564194 |
| SRR1048856 | 99.2 | 0.389823279 | 134 | -0.2964228 | 97201 | -0.347823837 | 0.368853595 | -0.342106941 | 0.296422761 | 0.342106941 | 0.680529144 |
| SRR1013538 | 91.7 | -0.052081905 | 448 | 0.6933547 | 19860 | -0.696635509 | 5.123438556 | 4.41247802 | -0.693354697 | -4.41247802 | -5.85455013 |
| SRR1013587 | 92.7 | 0.006838786 | 455 | 0.7154198 | 18809 | -0.70137557 | 3.819693455 | 3.108732919 | -0.715419799 | -3.108732919 | -4.518689501 |
| SRR1048846 | 99.1 | 0.38393121 | 124 | -0.3279443 | 100681 | -0.332128868 | 0.381100381 | -0.329860155 | 0.327944336 | 0.329860155 | 0.709606833 |
| SRR1003105 | 52.2 | -2.379449206 | 1396 | 3.68160002 | 2256 | -0.776030405 | 27.85706175 | 27.14610121 | -3.681600016 | -27.14610121 | -33.98318084 |
| SRR1048839 | 99.1 | 0.38393121 | 130 | -0.3090314 | 94976 | -0.357858695 | 0.384538322 | -0.326422215 | 0.309031391 | 0.326422215 | 0.66152612 |
| SRR1013661 | 84.1 | -0.499879158 | 614 | 1.21661284 | 12678 | -0.729026678 | 5.796514986 | 5.08555445 | -1.216612843 | -5.08555445 | -7.531073129 |
| SRR1013557 | 96 | 0.201277067 | 284 | 0.17640087 | 35227 | -0.627329591 | 2.471496933 | 1.760536397 | -0.176400865 | -1.760536397 | -2.362989786 |
| SRR1013591 | 89.1 | -0.205275702 | 493 | 0.83520178 | 17822 | -0.705826988 | 6.417344501 | 5.706383965 | -0.835201785 | -5.706383965 | -7.452688439 |
| SRR1013545 | 92.6 | 0.000946717 | 421 | 0.60824644 | 22170 | -0.686217296 | 3.214846119 | 2.503885582 | -0.608246444 | -2.503885582 | -3.797402605 |
| SRR1047983 | 98.5 | 0.348578795 | 179 | -0.1545757 | 51581 | -0.553572256 | 0.796457291 | 0.085496754 | 0.154575673 | -0.085496754 | -0.135914542 |

*Multiplying no. of contigs and unmapped reads with -1, as lower values show high quality genomes

| **Range of completeness parameters including complete single-copy, duplicated, fragmented, and missing BUSCO.** | | | | | |
| --- | --- | --- | --- | --- | --- |
| **Range** | **No. of Complete Single-copy BUSCO** | **Range** | **No. of Complete Duplicate BUSCO** | **No. of Fragmented BUSCO** | **No. of Missing BUSCO** |
| **>90%** | 392 | <2% | 444 | 370 | 372 |
| **80%-90%** | 45 | >2% | 30 | 104 | 102 |
| **<80%** | 37 |  |  |  |  |
| **Range of contiguity (Number of contigs, N50), and accuracy (unmapped reads) parameters.** | | | | | |
| **Range** | **N50** | **Range** | **No. of Contigs** | **Range** | **Unmapped Reads** |
| **>50 kb** | 354 | <200 | 346 | <2% | 348 |
| **<50 kb** | 120 | 200-500 | 69 | 2%-5% | 64 |
|  |  | >500 | 59 | >5% | 62 |

**Ranking of Genomes Based on the Z-score**

| **High-ranked (blue filled cells) and Low-ranked (Yellow filled) genomes** | |
| --- | --- |
| **SRA #** | **Summed z-score** |
| SRR7209071 | 10.6121165 |
| SRR10887300 | 10.6121165 |
| SRR9062699 | 5.756535399 |
| SRR10911065 | 5.364558057 |
| SRR7235686 | 5.364558057 |
| SRR10887303 | 4.917270772 |
| SRR7235685 | 4.917270772 |
| SRR11787122 | 3.881305327 |
| SRR7235688 | 3.540706324 |
| SRR10887302 | 3.540706324 |
| SRR11787123 | 3.098816229 |
| SRR3438086 | 3.0031397 |
| SRR11787114 | 2.97175263 |
| SRR11787129 | 2.961165678 |
| SRR11787110 | 2.948415686 |
| SRR6926203 | 2.936099354 |
| SRR11787125 | 2.931709992 |
| SRR7235666 | 2.862737477 |
| SRR10887299 | 2.862737477 |
| SRR11787130 | 2.862622924 |
| SRR11787121 | 2.848092331 |
| SRR4428965 | 2.835512373 |
| SRR10911062 | 2.822635346 |
| SRR11787112 | 2.820218538 |
| SRR11362441 | 2.774689663 |
| SRR11787128 | 2.761186616 |
| SRR11787111 | 2.753401066 |
| SRR11787126 | 2.710762429 |
| SRR7235684 | 2.672037988 |
| SRR10911063 | 2.672037988 |
| SRR11787127 | 2.663887823 |
| SRR7235689 | 2.652901004 |
| SRR3453146 | 2.628843199 |
| SRR5182484 | 2.611547935 |
| SRR11486365 | 2.603133802 |
| SRR1685377 | 2.582413172 |
| SRR1182722 | 2.57983017 |
| SRR11787116 | 2.500184608 |
| SRR7702518 | 2.499305267 |
| SRR6784331 | 2.458981384 |
| SRR11787115 | 2.431543656 |
| SRR6806305 | 2.424081412 |
| SRR11787117 | 2.4059987 |
| SRR1636908 | 2.388644025 |
| SRR7547846 | 2.35746826 |
| SRR11787118 | 2.243247055 |
| SRR6304499 | 2.214625715 |
| SRR11486381 | 2.037036873 |
| SRR5571307 | 2.012319473 |
| SRR7235878 | 1.996893235 |
| SRR10887301 | 1.996893235 |
| SRR5811620 | 1.980037728 |
| SRR6303974 | 1.961249288 |
| SRR1610005 | 1.946254882 |
| SRR7136124 | 1.940539353 |
| SRR2878463 | 1.856724931 |
| SRR3438087 | 1.853996632 |
| SRR11787108 | 1.819661167 |
| SRR7359081 | 1.8003559 |
| SRR7136076 | 1.776788123 |
| SRR11787109 | 1.743124621 |
| SRR2585813 | 1.736510584 |
| SRR7209078 | 1.720882216 |
| SRR10911067 | 1.720882216 |
| SRR7136074 | 1.688690177 |
| SRR7136105 | 1.644240763 |
| SRR7136122 | 1.611058926 |
| SRR8182744 | 1.573309786 |
| SRR7136080 | 1.562192906 |
| SRR2585418 | 1.545408534 |
| SRR11192678 | 1.5452567 |
| SRR7136062 | 1.541376413 |
| SRR7136035 | 1.540776797 |
| SRR7136125 | 1.535852785 |
| SRR7136030 | 1.509092371 |
| SRR3438067 | 1.508839654 |
| SRR7136119 | 1.505981772 |
| SRR7136050 | 1.503093537 |
| SRR7136057 | 1.502829316 |
| SRR7136098 | 1.454889869 |
| SRR7136097 | 1.446552235 |
| SRR7136093 | 1.441273112 |
| SRR7136103 | 1.437267406 |
| SRR7136095 | 1.430327698 |
| SRR7136049 | 1.424392343 |
| SRR7136126 | 1.418170928 |
| SRR7136048 | 1.416585711 |
| SRR1288379 | 1.404507349 |
| SRR7136099 | 1.393930387 |
| SRR5071100 | 1.392409415 |
| SRR7136052 | 1.388040498 |
| SRR7136111 | 1.374230306 |
| SRR7136104 | 1.361582887 |
| SRR7136032 | 1.359276227 |
| SRR7136066 | 1.358632865 |
| SRR7136039 | 1.355824808 |
| SRR7136063 | 1.342867018 |
| SRR11787120 | 1.302127486 |
| SRR7136042 | 1.300950646 |
| SRR7136045 | 1.299674788 |
| SRR7136046 | 1.2968489 |
| SRR7136121 | 1.291044943 |
| SRR7136088 | 1.287367218 |
| SRR7136041 | 1.28716724 |
| SRR7136037 | 1.286417341 |
| SRR7136106 | 1.283630579 |
| SRR7136038 | 1.273925609 |
| SRR7136123 | 1.263369197 |
| SRR1805599 | 1.26248405 |
| SRR7136040 | 1.256530034 |
| SRR7136047 | 1.234593584 |
| SRR7136115 | 1.233704777 |
| SRR7136113 | 1.232571737 |
| SRR1709626 | 1.23066209 |
| SRR7136078 | 1.228916457 |
| SRR7136109 | 1.222943111 |
| SRR7136108 | 1.222876654 |
| SRR7136073 | 1.207642702 |
| SRR7136053 | 1.200377926 |
| SRR7136065 | 1.196572963 |
| SRR7136112 | 1.188265176 |
| SRR7136060 | 1.18478653 |
| SRR7136100 | 1.182092422 |
| SRR7136091 | 1.178199324 |
| SRR7136096 | 1.178051955 |
| SRR7826315 | 1.176510906 |
| SRR7136084 | 1.175620712 |
| SRR1726160 | 1.173717736 |
| SRR7136077 | 1.170638501 |
| SRR7136064 | 1.160891841 |
| SRR7136033 | 1.159370931 |
| SRR7136086 | 1.156561001 |
| SRR7136107 | 1.150652493 |
| SRR1049066 | 1.150403895 |
| SRR7136036 | 1.144993116 |
| SRR2532675 | 1.134632072 |
| SRR7136094 | 1.119563539 |
| SRR7136081 | 1.10909144 |
| SRR3457705 | 1.109074674 |
| SRR1946930 | 1.101052871 |
| SRR7136092 | 1.096550961 |
| SRR7136120 | 1.095645059 |
| SRR1048843 | 1.09005955 |
| SRR7136079 | 1.083072185 |
| SRR7136044 | 1.076911004 |
| SRR7136043 | 1.066783877 |
| SRR7136085 | 1.061892139 |
| SRR1048842 | 1.054451417 |
| SRR7136058 | 1.037725489 |
| SRR7136034 | 1.034823223 |
| SRR1048882 | 1.029287761 |
| SRR7136089 | 1.028999749 |
| SRR2078207 | 1.024126188 |
| SRR1553800 | 1.024019591 |
| SRR1049074 | 1.02153012 |
| SRR7136101 | 1.01366297 |
| SRR7136117 | 1.008698402 |
| SRR1048855 | 0.998990428 |
| SRR7136029 | 0.998973864 |
| SRR1048880 | 0.995499625 |
| SRR2052522 | 0.993621327 |
| SRR7136061 | 0.976600497 |
| SRR1048854 | 0.972071467 |
| SRR1049077 | 0.967435413 |
| SRR7136069 | 0.964900846 |
| SRR1823703 | 0.964434186 |
| SRR7136102 | 0.96211615 |
| SRR7136082 | 0.955573043 |
| SRR1048857 | 0.945615538 |
| SRR1049067 | 0.942861962 |
| SRR1049061 | 0.942499916 |
| SRR1048858 | 0.934011914 |
| SRR1048978 | 0.931928275 |
| SRR1049052 | 0.925463304 |
| SRR1048979 | 0.923352581 |
| SRR7136031 | 0.920399843 |
| SRR1049075 | 0.9177065 |
| SRR1049009 | 0.910726674 |
| SRR1049002 | 0.910442906 |
| SRR1048973 | 0.909697476 |
| SRR1048881 | 0.908873885 |
| SRR1048977 | 0.900581995 |
| SRR1048851 | 0.897619732 |
| SRR4429027 | 0.890264903 |
| SRR1003106 | 0.886730381 |
| SRR1049055 | 0.884475667 |
| SRR3438088 | 0.88318585 |
| SRR1049059 | 0.879260038 |
| SRR1049057 | 0.878083361 |
| SRR1048850 | 0.876241554 |
| SRR1049056 | 0.875850838 |
| SRR1048971 | 0.873501531 |
| SRR1048845 | 0.857151472 |
| SRR1049064 | 0.855379109 |
| SRR7136087 | 0.852494416 |
| SRR1048863 | 0.847283339 |
| SRR1049053 | 0.8446102 |
| SRR1049049 | 0.839231639 |
| SRR1049072 | 0.837556811 |
| SRR1049033 | 0.834583473 |
| SRR7136059 | 0.832464564 |
| SRR1049070 | 0.830819034 |
| SRR1049020 | 0.830129705 |
| SRR1049065 | 0.828089216 |
| SRR1049036 | 0.824224658 |
| SRR1048970 | 0.820381209 |
| SRR1049010 | 0.818706493 |
| SRR1049019 | 0.817804152 |
| SRR1049045 | 0.814683519 |
| SRR1049069 | 0.814209272 |
| SRR7136114 | 0.80986493 |
| SRR1048859 | 0.804869923 |
| SRR1048841 | 0.802219774 |
| SRR7136071 | 0.795538796 |
| SRR7136118 | 0.785702811 |
| SRR1049025 | 0.783237376 |
| SRR1048962 | 0.769928795 |
| SRR1049038 | 0.767202522 |
| SRR1049073 | 0.757236413 |
| SRR1918970 | 0.749274274 |
| SRR1048999 | 0.74614216 |
| SRR1049044 | 0.745114898 |
| SRR1048879 | 0.741407985 |
| SRR1003101 | 0.738507255 |
| SRR1048836 | 0.735894688 |
| SRR1049048 | 0.732468019 |
| SRR1048864 | 0.72903688 |
| SRR1049037 | 0.722515567 |
| SRR1049063 | 0.717302609 |
| SRR7136067 | 0.716798485 |
| SRR1048835 | 0.716286634 |
| SRR1048972 | 0.714481996 |
| SRR1049000 | 0.709673502 |
| SRR1048846 | 0.70879817 |
| SRR1048844 | 0.699600849 |
| SRR1048963 | 0.699570367 |
| SRR1049029 | 0.690725889 |
| SRR10419414 | 0.684951179 |
| SRR1049060 | 0.682888972 |
| SRR1048838 | 0.681170203 |
| SRR1048856 | 0.679682594 |
| SRR1049050 | 0.661740708 |
| SRR1048840 | 0.661628088 |
| SRR1048839 | 0.660676598 |
| SRR1049001 | 0.659745327 |
| SRR1049071 | 0.649250757 |
| SRR7136051 | 0.646955203 |
| SRR1003102 | 0.644166214 |
| SRR7136110 | 0.642185216 |
| SRR1049051 | 0.64201491 |
| SRR1049032 | 0.634765446 |
| SRR2174092 | 0.630580389 |
| SRR9062702 | 0.629796051 |
| SRR1049047 | 0.619422281 |
| SRR1049068 | 0.608836339 |
| SRR1049028 | 0.60812397 |
| SRR7136075 | 0.607633686 |
| SRR6919274 | 0.602815539 |
| SRR1182726 | 0.600688281 |
| SRR10419390 | 0.59331864 |
| SRR1048847 | 0.586751964 |
| SRR1049076 | 0.580293333 |
| SRR1049046 | 0.571037379 |
| SRR1049058 | 0.546779782 |
| SRR1049054 | 0.533144314 |
| SRR1048976 | 0.507106444 |
| SRR1144817 | 0.490022921 |
| SRR2585419 | 0.486680486 |
| SRR1144768 | 0.467288607 |
| SRR1144774 | 0.453441202 |
| SRR10419565 | 0.43523725 |
| SRR1144807 | 0.43283423 |
| SRR1144725 | 0.431418863 |
| SRR1652540 | 0.428656926 |
| SRR1144796 | 0.420970456 |
| SRR7136068 | 0.418169332 |
| SRR1144733 | 0.392937341 |
| SRR1049043 | 0.37492466 |
| SRR1049035 | 0.373176773 |
| SRR1049042 | 0.370898075 |
| SRR11486432 | 0.35916593 |
| SRR1144757 | 0.350833767 |
| SRR7136070 | 0.348006137 |
| SRR1144787 | 0.327706667 |
| SRR1144805 | 0.31911677 |
| SRR1144790 | 0.296248279 |
| SRR7547837 | 0.29158429 |
| SRR1144813 | 0.288924243 |
| SRR5434330 | 0.283587496 |
| SRR2910877 | 0.281947907 |
| SRR4733931 | 0.28188277 |
| SRR1144820 | 0.280269538 |
| SRR1144776 | 0.279535989 |
| SRR1144749 | 0.257509034 |
| SRR1049034 | 0.256450622 |
| SRR1503321 | 0.234152343 |
| SRR1144754 | 0.208151556 |
| SRR1144780 | 0.167221019 |
| SRR1144771 | 0.146746712 |
| SRR11589263 | 0.126145296 |
| SRR1048871 | 0.096273176 |
| SRR1144766 | 0.092109354 |
| SRR4733932 | 0.058259848 |
| SRR1048872 | 0.040092562 |
| SRR5434334 | -0.009807526 |
| SRR1049024 | -0.020138921 |
| SRR10419106 | -0.068619034 |
| SRR7136072 | -0.12597145 |
| SRR1047983 | -0.137116209 |
| SRR7136056 | -0.143025023 |
| SRR2124318 | -0.165235358 |
| SRR1048990 | -0.411981517 |
| SRR11648067 | -0.413923405 |
| SRR2174058 | -0.419586112 |
| SRR1144783 | -0.502481879 |
| SRR10911066 | -0.547101943 |
| SRR1048989 | -0.569654281 |
| SRR1652534 | -0.572730638 |
| SRR1805641 | -0.662532492 |
| SRR5434331 | -0.673932149 |
| SRR7235687 | -0.838378715 |
| SRR7136054 | -0.843892495 |
| SRR4429026 | -0.962062055 |
| SRR6921434 | -1.161772774 |
| SRR1048010 | -1.212127956 |
| SRR7136055 | -1.226148912 |
| SRR1049062 | -1.23585743 |
| SRR1047988 | -1.275732689 |
| SRR1144737 | -1.286866163 |
| SRR1049039 | -1.348247941 |
| SRR1816847 | -1.375657839 |
| SRR1013541 | -1.544486089 |
| SRR1013576 | -1.740157088 |
| SRR1013571 | -1.752180956 |
| SRR1050530 | -1.870744014 |
| SRR1363336 | -1.884426545 |
| SRR1013555 | -1.92750978 |
| SRR1013575 | -1.974033385 |
| SRR1013670 | -2.070524033 |
| SRR1013572 | -2.075990947 |
| SRR1013556 | -2.107287086 |
| SRR1013574 | -2.145961166 |
| SRR1013639 | -2.147187041 |
| SRR1013640 | -2.194974247 |
| SRR1013558 | -2.274383341 |
| SRR1803044 | -2.326779595 |
| SRR1013557 | -2.364693213 |
| SRR9853553 | -2.381486647 |
| SRR1013641 | -2.396565776 |
| SRR1013559 | -2.483212631 |
| SRR1013638 | -2.538973278 |
| SRR1013573 | -2.765958862 |
| SRR1013543 | -2.934480218 |
| SRR1049004 | -3.083721229 |
| SRR1048865 | -3.106162227 |
| SRR1013606 | -3.129918439 |
| SRR1013542 | -3.132676136 |
| SRR1048866 | -3.137886185 |
| SRR1013539 | -3.248996562 |
| SRR1048867 | -3.287423863 |
| SRR11648057 | -3.329798043 |
| SRR1048997 | -3.368722525 |
| SRR1049005 | -3.36991059 |
| SRR1047984 | -3.375991636 |
| SRR1561272 | -3.384540425 |
| SRR1013642 | -3.400371841 |
| SRR1048986 | -3.442146737 |
| SRR1048868 | -3.447884173 |
| SRR1048985 | -3.496295148 |
| SRR1013624 | -3.53507602 |
| SRR1013643 | -3.626815725 |
| SRR1048998 | -3.633788928 |
| SRR1013544 | -3.647777874 |
| SRR1013619 | -3.682401359 |
| SRR10911064 | -3.726639277 |
| SRR1013545 | -3.799733752 |
| SRR1013546 | -3.819578697 |
| SRR1013605 | -3.890753041 |
| SRR1048981 | -3.902767832 |
| SRR1048980 | -3.921749762 |
| SRR1013653 | -3.930353404 |
| SRR1013551 | -3.985329118 |
| SRR1013540 | -4.009257664 |
| SRR1013560 | -4.032348332 |
| SRR1013552 | -4.09425332 |
| SRR1013620 | -4.097361833 |
| SRR1048995 | -4.445892865 |
| SRR1048996 | -4.457010667 |
| SRR1048861 | -4.473095919 |
| SRR1013657 | -4.51084036 |
| SRR1013587 | -4.5211271 |
| SRR1013652 | -4.690422453 |
| SRR1013594 | -4.805450334 |
| SRR1013656 | -4.829769408 |
| SRR1013650 | -5.230576559 |
| SRR1013586 | -5.415355585 |
| SRR1013588 | -5.415727084 |
| SRR1013651 | -5.436653227 |
| SRR1013623 | -5.465096221 |
| SRR1013593 | -5.642133175 |
| SRR1013592 | -5.652911465 |
| SRR1013666 | -5.758753493 |
| SRR1013680 | -5.783438233 |
| SRR1013649 | -5.784401291 |
| SRR1013538 | -5.857016039 |
| SRR1013537 | -5.905161561 |
| SRR1013667 | -5.939081054 |
| SRR1013655 | -5.995876679 |
| SRR1048862 | -6.009930657 |
| SRR1013627 | -6.079850149 |
| SRR1013564 | -6.248075636 |
| SRR1013677 | -6.339689157 |
| SRR1013648 | -6.360907756 |
| SRR1013585 | -6.424580062 |
| SRR1013679 | -6.499559723 |
| SRR1013628 | -6.512535001 |
| SRR1013616 | -6.580014009 |
| SRR1013633 | -6.843566797 |
| SRR1013579 | -6.872739723 |
| SRR1013654 | -6.961420575 |
| SRR1013570 | -6.981743505 |
| SRR1013569 | -7.037917948 |
| SRR1013615 | -7.043236371 |
| SRR1013547 | -7.047308084 |
| SRR1013658 | -7.0761017 |
| SRR1013683 | -7.10995973 |
| SRR1013676 | -7.13226337 |
| SRR1013659 | -7.14857706 |
| SRR1013580 | -7.199378667 |
| SRR1013563 | -7.202304127 |
| SRR1013645 | -7.238865144 |
| SRR1013671 | -7.239856911 |
| SRR1013662 | -7.242026442 |
| SRR1013631 | -7.26158904 |
| SRR1013665 | -7.311149789 |
| SRR1013634 | -7.311944572 |
| SRR1013644 | -7.373201359 |
| SRR1013625 | -7.394639647 |
| SRR1013568 | -7.4442609 |
| SRR1013591 | -7.455429536 |
| SRR1013668 | -7.50842259 |
| SRR1013661 | -7.534447658 |
| SRR1013664 | -7.554816221 |
| SRR1013673 | -7.565818349 |
| SRR1013626 | -7.678527686 |
| SRR1013632 | -7.698986844 |
| SRR1730387 | -7.722572225 |
| SRR1013601 | -7.731241673 |
| SRR1013660 | -7.901716719 |
| SRR1013602 | -8.016347588 |
| SRR1013561 | -8.142415008 |
| SRR1013562 | -8.147782877 |
| SRR1013669 | -8.213672314 |
| SRR1013663 | -8.281013272 |
| SRR1013682 | -8.375433277 |
| SRR1509623 | -8.447699087 |
| SRR1013672 | -8.993478941 |
| SRR1013567 | -9.411099938 |
| SRR1013646 | -10.30164086 |
| SRR1013596 | -10.49317785 |
| SRR1003103 | -10.50735171 |
| SRR1013647 | -10.76811393 |
| SRR1013595 | -10.90958966 |
| SRR1013674 | -12.89164281 |
| SRR1013675 | -13.68101169 |
| SRR1013583 | -13.82590332 |
| SRR1013584 | -13.92086709 |
| SRR1013613 | -14.81127991 |
| SRR1013614 | -14.84171366 |
| SRR1013566 | -16.06830453 |
| SRR1013612 | -16.97703403 |
| SRR1013611 | -17.61129995 |
| SRR1013565 | -18.48587869 |
| SRR1003105 | -33.99053287 |
